# Connexin-43 links Neuromesodermal progenitor states to segmentation clock robustness during vertebrate axis elongation

**DOI:** 10.64898/2026.08.27.747369

**Authors:** Jonas Cruzel, Romane Bertrand, Felipe Maurelia, Prikshit, Enola Ranvier, Charlène Guillot

## Abstract

Neuromesodermal progenitors (NMPs) sustain vertebrate body-axis elongation by generating both neural and paraxial mesodermal tissues. Although signaling and metabolic pathways regulate NMP states, whether intercellular communication contributes to the coordination of progenitor behaviour and developmental timing remains unclear. Here, we identify GJA1, encoding connexin-43 (Cx43), as a gene dynamically enriched within the neuromesodermal competent domain of the chick embryo. Cx43-associated channels and hemichannels accumulate preferentially within the NMP population, and a photoactivatable tracer assay demonstrates enhanced connexin-mediated exchange within the posterior growth zone. Pharmacological inhibition of hemichannels or gap junctions revealed distinct contributions of these communication modes to transcriptional regulation across the NMP continuum. Gap-junction inhibition primarily altered SOX2 expression within progenitor populations, whereas hemichannel inhibition selectively affected TBXT expression in mesodermal cells. Connexin inhibition also reduced the relative size of the progenitor compartment and altered the spatial organization of newly formed somites. Strikingly, disruption of connexin-mediated communication impaired segmentation dynamics, leading to increased frequencies of off-pace segmentation events, accelerated segmentation timing and progressive deviation from the expected segmentation program. These defects emerged rapidly and accumulated over successive segmentation cycles, indicating a requirement for connexin activity in maintaining developmental robustness. Together, our findings identify connexin-43 as a regulator of neuromesodermal progenitor states and reveal a previously unrecognized link between intercellular communication and segmentation clock robustness during vertebrate axis elongation.

## INTRODUCTION

The elongation of the vertebrate body axis depends on the coordinated activity of neuromesodermal progenitors (NMPs), a population of bipotent cells located in the posterior growth zone, also referred to as the neuromesodermal competent (NMC) domain, that contribute to both neural and paraxial mesodermal lineages. The NMP population within the NMC domain is characterized by the co-expression of SOX2 and TBXT, and their maintenance relies on a dynamic balance between neural and mesodermal transcriptional programs controlled by WNT, FGF and retinoic acid signalling (Gouti et al., 2014; Henrique et al., 2015; Koch et al., 2017; Steventon and Martinez Arias, 2017; Binagui-Casas et al., 2021; Wymeersch et al., 2021). A more recent work using pluripotent stem cells, including human induced pluripotent stem cell–derived NMP-like cells, has refined this model and it suggests that NMPs comprise a continuum of transcriptional states that dynamically respond to environmental and signaling inputs (Tsakiridis and Wilson, 2015; Attardi et al., 2018; Frith et al., 2018; Verrier et al., 2018).

Beyond these signaling pathways, metabolic regulation has emerged as a critical component of posterior axis development. In particular, glycolytic activity has been shown to influence both NMC cell identity and segmentation dynamics. Perturbations of glucose metabolism alter the balance between neural and mesodermal programs and modulate segmentation clock activity, linking metabolic state to both cell fate and embryonic patterning (Oginuma et al., 2017; Oginuma et al., 2020). Recent work using human *in vitro* segmentation models has further demonstrated a direct coupling between metabolic rate and segmentation clock dynamics, showing that changes in cellular metabolism can control oscillatory periodicity and timing (Dias-Cuadros et al., 2020; Dias-Cuadros et al., 2023). Together, these results indicate that metabolic state directly influences progenitor states and segmentation timing. Consistent with this regulatory complexity, our previous single-cell transcriptomic analysis of the posterior domain (Guillot et al., 2021) identified GJA1 as a core gene enriched in the NMP cluster, raising the question of whether connexin-mediated communication contributes to the coordination of NMPs states and segmentation dynamics.

At the tissue level, the conversion of paraxial mesoderm into somites is controlled by the segmentation clock, a molecular oscillator that ensures the periodic formation of somites with remarkable precision. In amniotes, somites form at defined intervals (approximately 90 minutes in the chick embryo), reflecting a highly robust oscillatory system. While modulation of signaling pathways or environmental parameters can alter the speed of the segmentation clock, such perturbations typically lead to uniform acceleration or deceleration while preserving periodicity. In developmental systems, robustness refers to the capacity to generate reproducible outcomes despite intrinsic sources of biological noise, including stochastic variation in gene expression, cell behavior and environmental fluctuations (Oates et al., 2012; Sonnen et al., 2018). In contrast, irregular segmentation, characterized by variable intervals, pauses, or the formation of multiple somites in rapid succession, is comparatively rare and is generally associated with defects in synchronization between oscillating cells (Dequeant and Pourquié, 2008; Oates et al., 2012; Sonnen et al., 2018; Meijer and Sonnen, 2024). Together, these observations suggest that mechanisms coordinating cell behavior at the tissue scale are essential for maintaining segmentation precision (Oates et al., 2012; Sonnen et al., 2018; Hubaud et al., 2017). However, the processes ensuring such coordination remain incompletely understood.

Mechanisms responsible for coordinating cellular behaviour across the posterior domain are still poorly defined. Connexins, which form gap junctions and hemichannels, represent strong candidates for mediating such coupling, as they enable the exchange of ions and small signalling molecules between cells and with the extracellular environment. Connexin-43 (Cx43), encoded by GJA1, is one of the most widely expressed connexins and plays essential roles in tissue morphogenesis and physiological coordination. In the developing heart, connexin-mediated coupling is required for the propagation of electrical signals and the establishment of rhythmic contractions, illustrating how intercellular communication regulates dynamic and oscillatory behaviors at the tissue scale (Evans and Martin, 2002; Goodenough and Paul, 2003). Importantly, connexins can operate through two distinct modes of communication: gap junctions, which directly connect adjacent cells, and hemichannels, which open to the extracellular space and regulate the release or uptake of signaling molecules. Gap junctions mediate direct cell–cell coupling, whereas hemichannels are thought to regulate the extracellular signaling environment.

However, whether connexin-mediated communication contributes to the coordination of NMP states and segmentation dynamics remains unclear. In our previous work (Guillot et al., 2021), we characterized NMP dynamics during chick axis elongation, highlighting the importance of coordinated cell behaviors within the posterior growth zone. Here, we address this question and show that GJA1 expression is dynamically enriched within the neuromesodermal competent domain and that connexin-43 channels are functionally active in these cells. We further demonstrate that pharmacological perturbation of connexin activity, using inhibitors targeting hemichannels and gap junctions, differentially affects transcriptional states within the NMP continuum, posterior tissue organization, and segmentation dynamics. Together, these results identify connexin-43 as a regulator of neuromesodermal progenitor states and segmentation clock robustness during vertebrate axis formation.

## RESULTS

### GJA1 expression is specifically enriched in the neuromesodermal competent domain

To determine whether connexin-mediated communication may contribute to posterior axis development, we first examined the spatial distribution of GJA1 expression. Previous analysis of single-cell RNA-sequencing data identified GJA1 among a restricted set of genes enriched within the NMP cluster (Guillot et al, 2021). We validated these observations *in situ* using multiplex HCR-FISH for GJA1, SOX2 and TBXT probes across multiple developmental stages (Fig. 1A–I). At HH7/HH8 (approximately 30 h of development), GJA1 expression was strongly enriched within the NMC domain, where SOX2 and TBXT expression overlap (Fig. 1A). Quantification of fluorescence intensity along the posterior axis revealed a distinct peak of GJA1 expression centered on the NMC region (Fig. 1D). At HH10/HH12 (approximately 48 h), GJA1 expression remained elevated within the NMC domain but also extended into the adjacent neural tissue, resulting in a broader expression profile across the posterior region (Fig. 1B,E). By HH18/HH20 (approximately 72 h), GJA1 expression was markedly reduced and displayed a more homogeneous distribution across the analyzed posterior tissues (Fig. 1C,F).

**Figure 1.**
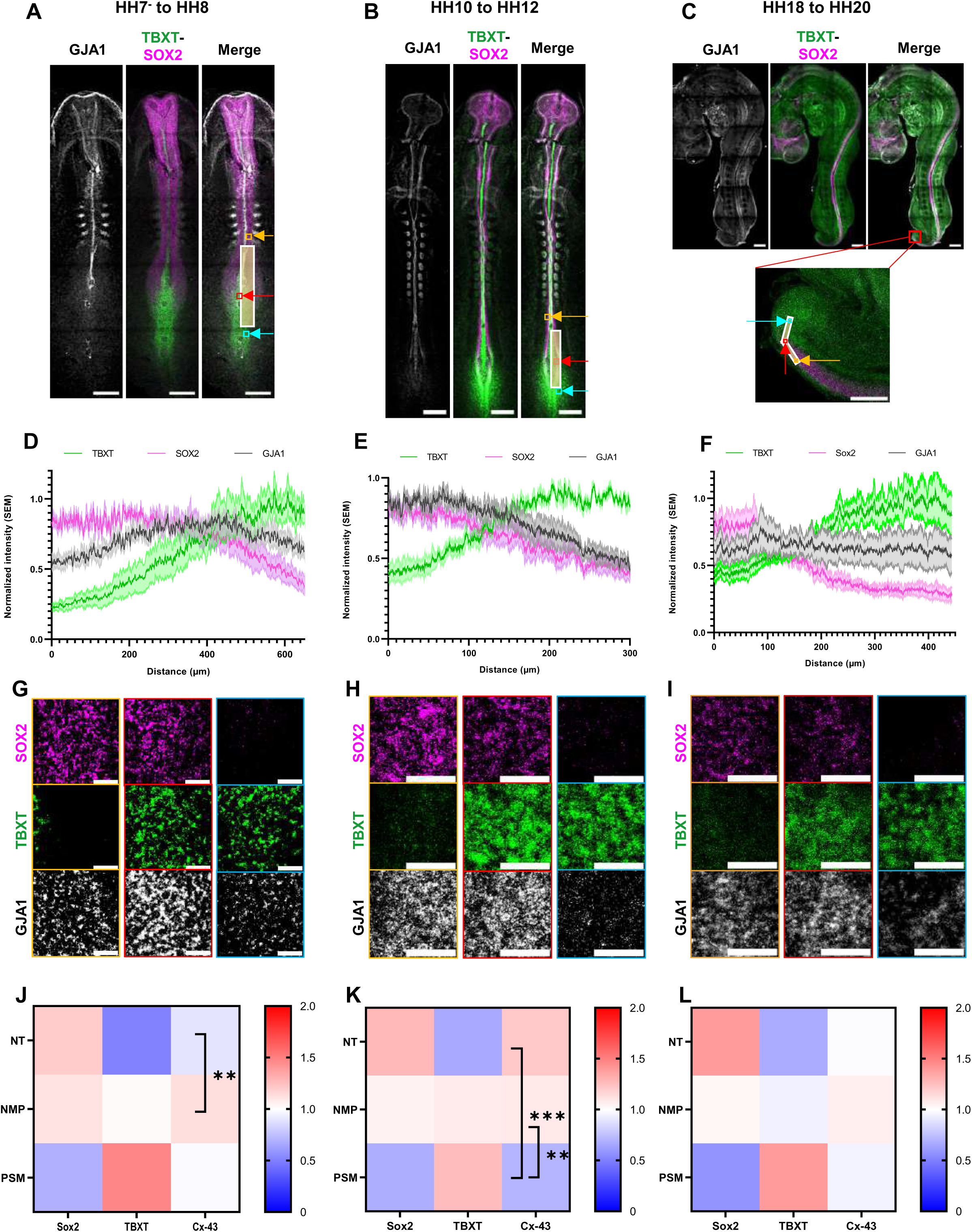
- GJA1 is dynamically enriched within the neuromesodermal competent domain during axis elongation. A–C, Representative embryos at HH7^-^–8, HH10–12 and HH18–20, respectively, hybridized with RNA probes targeting *GJA1*, *SOX2* and *TBXT* and imaged by confocal microscopy using ×10 and ×20 objectives. Scale bars, 300 μm for HH7^-^–8 and HH10–12; 500 μm for HH18–20; 200 μm for enlarged views. Lines overlaid on the embryos indicate the trajectories used for spatial quantification of probe intensity, extending from the neural tube through the neuromesodermal progenitor (NMP) domain to the presomitic mesoderm (PSM). Coloured boxes indicate the regions used for local intensity quantification. **D–F,** Normalized probe intensities plotted as a function of distance along the indicated trajectory, from the neural tube through the NMP domain to the presomitic mesoderm (PSM), at HH7-–8, HH10–12 and HH18–20, respectively. *SOX2*, pink; *TBXT*, green; *GJA1*, black. Shaded areas represent the standard error of the mean (s.e.m.) at each position along the trajectory. n = 6, 5 and 4 embryos for HH7-–8, HH10–12 and HH18–20, respectively. **G–I,** Representative 100 × 100 μm regions at HH7-–8 and 50 × 50 μm regions at HH10–12 and HH18– 20 showing probe expression in the neural tube (orange), NMP domain (red) and presomitic mesoderm (PSM; blue). Scale bars, 25 μm. n = 6, 5 and 4 embryos for HH7-–8, HH10–12 and HH18–20, respectively. **J–L,** Representative heatmaps showing normalized relative *GJA1* expression in the indicated tissues at HH7-–8 and HH10–12; 500 μm for HH18–20, respectively. Statistical significance between tissue pairs was assessed using pairwise unpaired two-tailed t-tests. At HH7-–8, *GJA1*expression was significantly different between the neural tube and NMPs (P = 0.00716). At HH10–12, significant differences were observed between PSM and NMPs (P = 0.000490) and between the neural tube and PSM (P = 0.00936). No significant differences were detected between tissue pairs at HH18–20. P values are unadjusted for multiple comparisons. n = 6, 5 and 4 embryos for HH7-–8, HH10–12 and HH18–20, respectively.

Thus, although GJA1 transcripts were detected throughout the posterior axis, expression was not uniform and displayed a clear enrichment within the NMC domain during active axis elongation. Specific quantification of transcripts in the NMP, Neural and PSM domains (Fig 1G-L) showed increased GJA1 expression in the NMPs at HH7/HH8 and HH10/HH12. Thus, cells occupying the core of the SOX2⁺/TBXT⁺ territory exhibited higher GJA1 expression than neighboring neural (SOX2-high/TBXT-low) and mesodermal (TBXT-high/SOX2-low) populations. Importantly, the spatial distribution of GJA1 evolved over developmental time. Expression was initially concentrated within the NMC domain, progressively broadened during intermediate stages, and became more homogeneous at later stages of development (Fig. 1J–L).

Together, these results demonstrate that GJA1 expression is dynamically enriched within the neuromesodermal competent domain during posterior axis elongation, raising the possibility that connexin-mediated communication contributes to progenitor function and tissue coordination in the posterior growth zone (Goodenough & Paul, 2003; Solan & Lampe, 2009).

### Cx43 channels and hemichannels are enriched within the neuromesodermal progenitor population

To determine whether transcriptional enrichment of GJA1 translates into increased formation of connexin-mediated communication structures, we analyzed the distribution of Cx43 puncta by immunostaining and image-based segmentation (Fig. 2A–G). To compare the distribution of these structures across posterior tissues, images were collected in the neural tube (NT), NMP, or presomitic mesoderm (PSM) domains (Fig. 2A). In order to separate noisy points from real Cx43 puncta, we built an ilastik project that allows to identify punctas in all the images. The mean intensity of the Cx43 puncta segmented over its background (Supp Data Fig.2A) indicates that the punctas are successfully identify even when images have different levels of background. As expected, Cx43 signal was detected as discrete punctate structures at the cell membrane, consistent with the presence of connexin channels and hemichannels (Fig. 2B and Supp Data Fig.2B).

**Figure 2.**
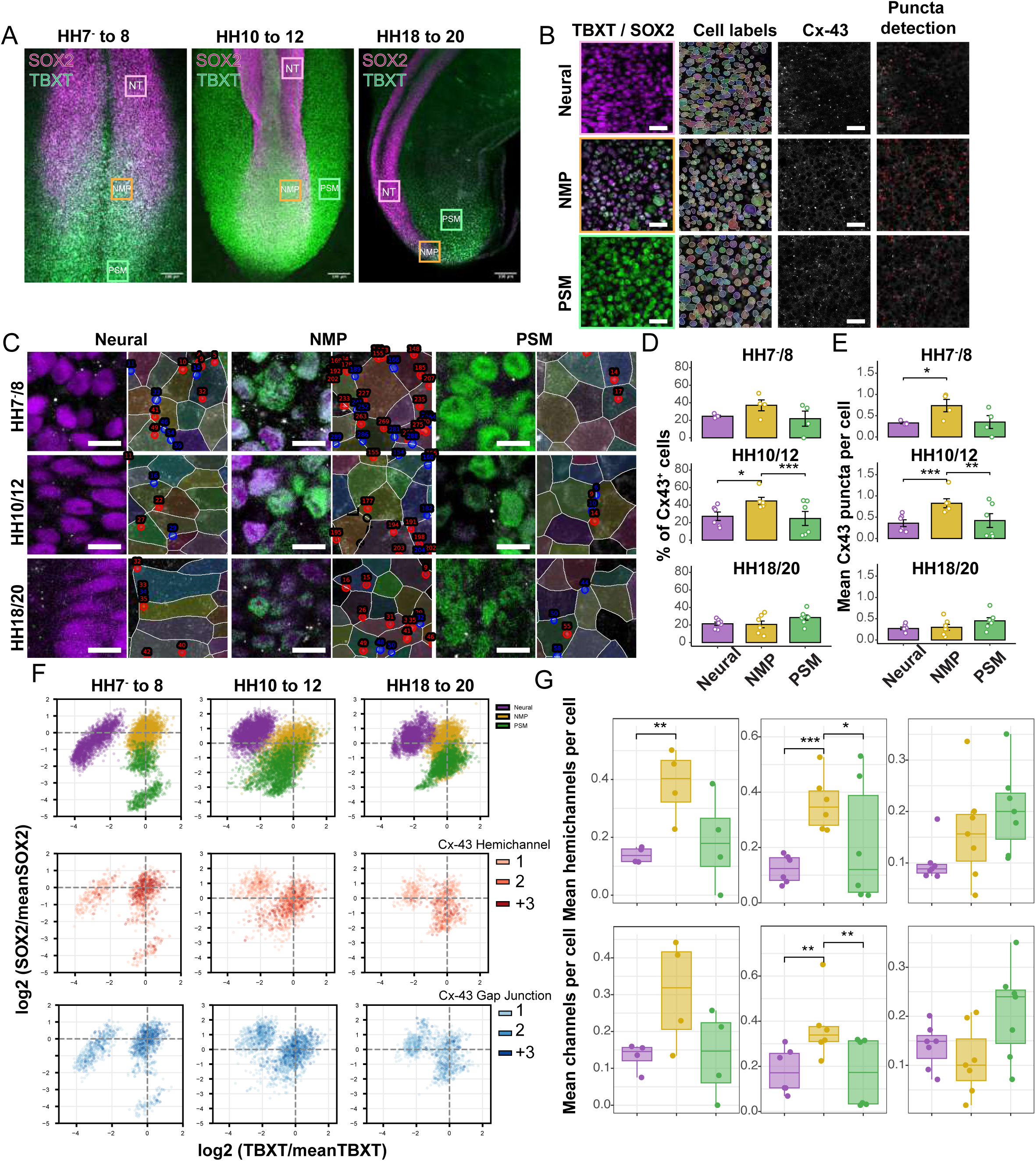
Neuromesodermal progenitors are enriched in Cx43 channels and hemichannels. **A**, Experimental workflow for quantification of Cx43 puncta. Posterior embryonic tissues were immunostained for SOX2, TBXT and Cx43, and images were acquired within the neural tube (NT), neuromesodermal progenitor (NMP) and presomitic mesoderm (PSM) domains. **B,** Representative confocal images showing SOX2, TBXT and Cx43 staining in the indicated tissue domains at HH7–8, HH10–12 and HH18–20. Cx43 puncta are detected at the cell membrane. **C,** Density maps showing the distribution of Cx43 hemichannel-associated puncta (red) and gap-junction-associated puncta (blue) projected onto the SOX2/TBXT continuum. Both classes of puncta are enriched within the central SOX2⁺/TBXT⁺ region corresponding to the NMP population. **D**, Quantification of the percentage of Cx43-positive cells within the NT, NMP and PSM populations at HH7–8, HH10–12 and HH18–20. NMPs exhibit a significantly higher proportion of Cx43-positive cells than neighboring neural and mesodermal tissues during active axis elongation. **E,** Quantification of the mean number of Cx43 puncta per cell within the NT, NMP and PSM populations at HH7–8, HH10–12 and HH18–20. NMPs display significantly higher puncta abundance than neighboring tissues at early developmental stages. **F,** Quantification of the mean number of hemichannel-associated puncta per cell within the NT, NMP and PSM populations at HH7–8, HH10–12 and HH18–20. **G,** Quantification of the mean number of gap-junction-associated puncta per cell within the NT, NMP and PSM populations at HH7–8, HH10–12 and HH18–20. Statistical significance was assessed using the Kruskal–Wallis test followed by Dunn’s multiple-comparisons test. Significant comparisons are indicated on the graphs. Data are presented as individual embryos with mean ± s.e.m. n = 6, 5 and 4 embryos for HH7–8, HH10–12 and HH18–20, respectively. Scale bars, 100μm.

After Cx43 assignation to individual cells segmented with cellprofiler, quantification revealed that Cx43-positive cells were not uniformly distributed across the different tissues. At both stages HH7/8 and HH10/12 of development, the NMP population exhibited a significantly higher proportion of Cx43-positive cells than neighboring neural and mesodermal populations (Fig. 2D). By contrast, these differences were largely lost by 72 h, when the distribution of Cx43-positive cells became more homogeneous across tissues. The same pattern is observed when we analyze the mean abundance of Cx43 puncta within individual cells. At 30 h and 48 h, NMPs contained significantly more Cx43 puncta per cell than neural tube and PSM populations (Fig. 2E), indicating that there are not just more cells positives for Cx43 puncta, but there is also more Cx43 puncta detected per cell. This enrichment progressively decreased at later developmental stages, mirroring the temporal dynamics observed for GJA1 expression.

Because connexin-43 can mediate communication through both hemichannels and gap-junction channels, we next examined the distribution of these two classes of structures across the SOX2/TBXT continuum. Density maps revealed that both hemichannel-associated (red) and channel-associated (blue) puncta were preferentially enriched within the central SOX2⁺/TBXT⁺ region corresponding to the NMP population (Fig. 2C,F). Quantitative analysis confirmed this enrichment. NMPs exhibited significantly higher densities of both hemichannel-associated and gap-junction-associated puncta than neighboring neural and mesodermal populations during active axis elongation (Fig. 2G). This enrichment was most pronounced at 30 h and 48 h and became less evident by 72 h, indicating that both forms of connexin-mediated communication are dynamically regulated during posterior development.

Together, these observations demonstrate that neuromesodermal progenitors are enriched not only in GJA1 expression and total Cx43 puncta, but also in both major forms of connexin-mediated communication. The preferential accumulation of hemichannels and gap-junction-associated channels within NMPs identifies this population as a highly communication-competent compartment of the posterior growth zone and suggests that connexin-mediated communication contributes to the regulation of progenitor behavior during axis elongation (Ton & Iovine, 2013; Yang et al., 2019).

### Connexin channels are functionally active within the NMC domain

To determine whether connexin channels are functionally active within the posterior axis, we employed a photoactivatable coumarin-based tracer assay in distinct posterior tissues (Fig. 3A,B). Local photoactivation induced a progressive decrease in fluorescence intensity within the activated region over time. Notably, the rate of fluorescence loss varied between tissues. Quantification of fluorescence intensity within both the photoactivated region and the surrounding non-photoactivated cells revealed that signal loss from the activated area was not accompanied by detectable accumulation of fluorescence in neighboring cells (Fig. 3C–F). These observations suggest that the tracer is released from the activated domain rather than extensively transferred between adjacent cells and are therefore consistent with connexin-mediated exchange with the extracellular environment.

**Figure 3.**
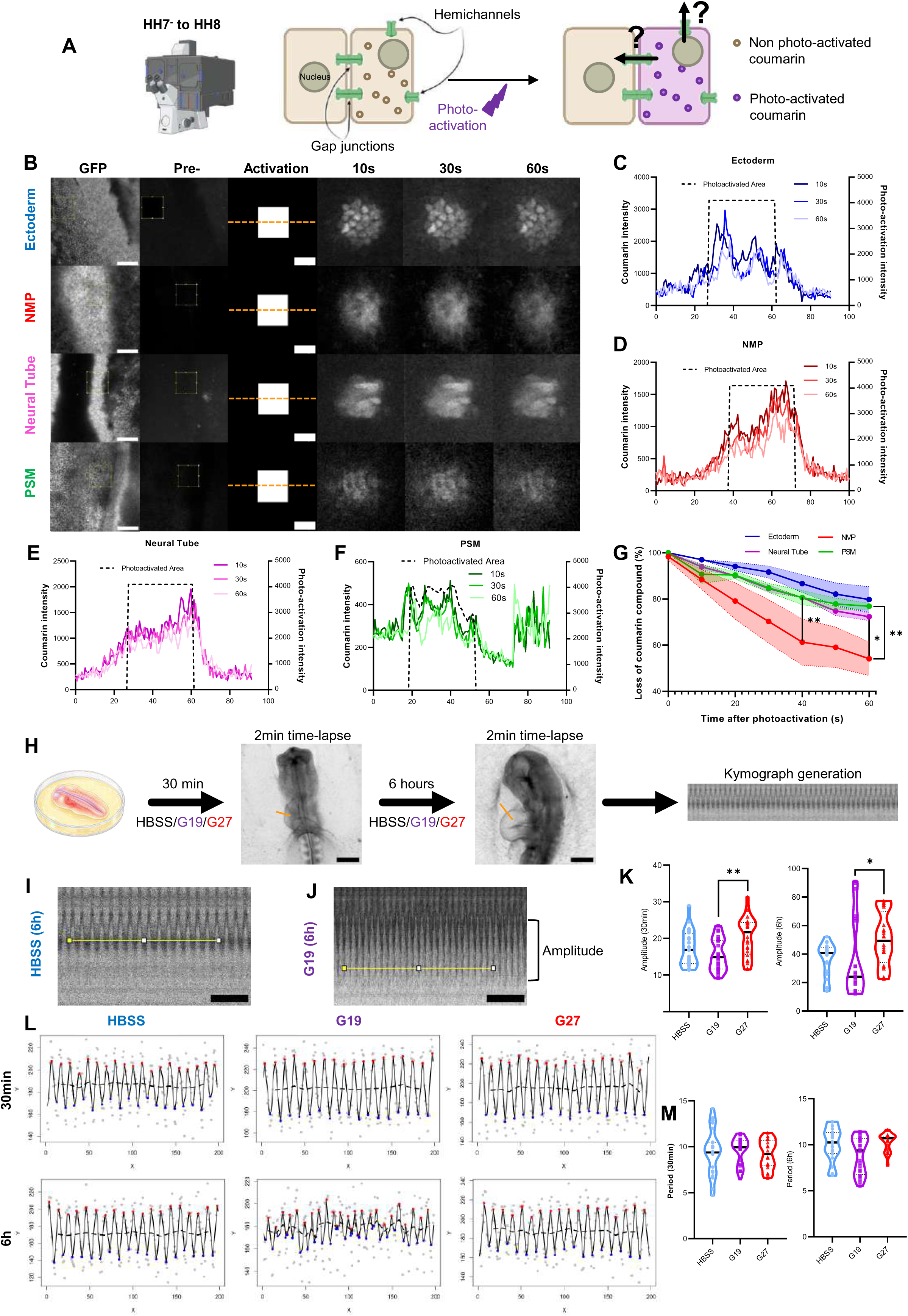
- Cx43-mediated communication is functionally active within the NMC domain and is efficiently inhibited by GAP19 and GAP27 A, Experimental design used to assess Cx-43 channel activity using the photoactivatable compound NPE-HCCC2/AM (coumarin). **B,** Representative images showing the selected imaging regions in the indicated tissues (yellow box; GFP and pre-activation images; scale bar, 100 μm), the photoactivation region (white box; scale bar, 10 μm) and time-lapse monitoring of fluorescence over 60 s. The 100-μm orange line indicates the trajectory used to quantify fluorescence loss over time. **C–F,** Representative fluorescence intensity profiles of photoactivatable coumarin over time, with successive time points shown from dark to light. The photoactivated region is indicated by the dotted line. **G,** Quantification of coumarin fluorescence loss over time in the indicated cell populations: ectoderm (blue), NMPs (red), neural tube (pink) and presomitic mesoderm (PSM; green). Statistical significance was assessed using pairwise two-sample Kolmogorov–Smirnov tests at the indicated time points. At 60 s, significant differences were observed between NMPs and PSM (P = 0.00820) and between NMPs and the neural tube (P = 0.0326); at 40 s, a significant difference was observed between NMPs and the neural tube (P = 0.00470). P values are unadjusted for multiple comparisons. n = 4 embryos. **H,** Experimental design used to assess the effects of Gap19 and Gap27 inhibition on embryonic heart function after 30 min and 6 h of treatment. Cardiac activity was assessed by kymography from 2-min time-lapse recordings. Scale bar, 500 μm. **I,J,** Representative kymographs after 6 h of treatment with HBSS and Gap19 (G19), respectively. The cardiac period (horizontal yellow line) and cardiac amplitude (black bracket) used for quantification are indicated. Scale bar, 2 s. **K,** Quantification of cardiac amplitude after 30 min and 6 h of treatment. Statistical significance was assessed using the Kruskal–Wallis test followed by Dunn’s multiple-comparisons test with adjusted P values. The Kruskal–Wallis test indicated significant differences among treatment groups at both 30 min (P = 0.00730) and 6 h (P = 0.0104). At 30 min, Dunn’s multiple-comparisons test showed no significant difference between HBSS and G19 or G27, but significant differences between G19 and G27 (P = 0.00590). At 6 h, no significant difference was observed between HBSS and G19 or G27, whereas significant differences were observed between G19 and G27 (P = 0.01060). n = 8 embryos per treatment group. **L,** Representative kymographs showing embryonic cardiac activity as a function of time and treatment condition **M,** Quantification of cardiac period after 30 min and 6 h of treatment. Statistical significance was assessed using the Kruskal–Wallis test followed by Dunn’s multiple-comparisons test with adjusted P values. The Kruskal–Wallis test showed no significant difference among treatment groups after 30 min, whereas a significant difference was observed after 6 h (P = 0.0330). At 6 h, Dunn’s multiple-comparisons test showed no significant differences between any groups. n = 8 embryos per treatment group.

Quantification of fluorescence decay revealed marked regional differences in tracer loss (Fig. 3G). The NMC domain exhibited significantly faster coumarin loss than the ectoderm, neural tube and presomitic mesoderm, indicating that connexin channels within the NMC are not only present but display enhanced functional activity. These observations identify the NMC as a region of elevated connexin-mediated exchange and suggest that communication through connexin channels is particularly active within the progenitor population.

To validate the pharmacological tools used throughout this study, we next assessed embryonic cardiac activity, a well-established physiological readout of connexin-dependent tissue coordination (Fig. 3H– M). During heart development, coordinated propagation of electrical activity relies on Cx43-mediated intercellular communication, making cardiac contraction dynamics a sensitive indicator of connexin function (Reaume et al., 1995; Ewart et al., 1997) We tested two inhibitors: GAP19, which preferentially blocks Cx43 hemichannels (Abudara et al, 2014), and GAP27, which preferentially inhibits gap-junction communication (Wang et al., 2013). Both treatments significantly altered cardiac contraction dynamics. Changes in contraction amplitude were detected after both 30 min and 6 h of treatment (Fig. 3I–K), whereas contraction periodicity was significantly affected after 6 h (Fig. 3L,M). These observations demonstrate that connexin-dependent communication is efficiently perturbed under our experimental conditions and that the effects of both inhibitors are detectable within 30 min and persist for at least 6 h.

Taken together, these findings demonstrate that connexin channels are functionally active within the NMC domain, validate GAP19 and GAP27 as effective pharmacological tools in the chick embryo, and establish that the developmental phenotypes observed in the posterior axis arise under conditions of impaired connexin-mediated communication.

### Connexin inhibition differentially affects SOX2 and TBXT expression across the NMP continuum

To determine whether connexin activity contributes to the regulation of NMP-associated transcriptional programmes, we quantified SOX2 and TBXT expression following pharmacological inhibition of connexin function and classified cells along the neural–mesodermal continuum using SOX2/TBXT density plots and single-cell segmentation analysis (Fig. 4A).

**Figure 4.**
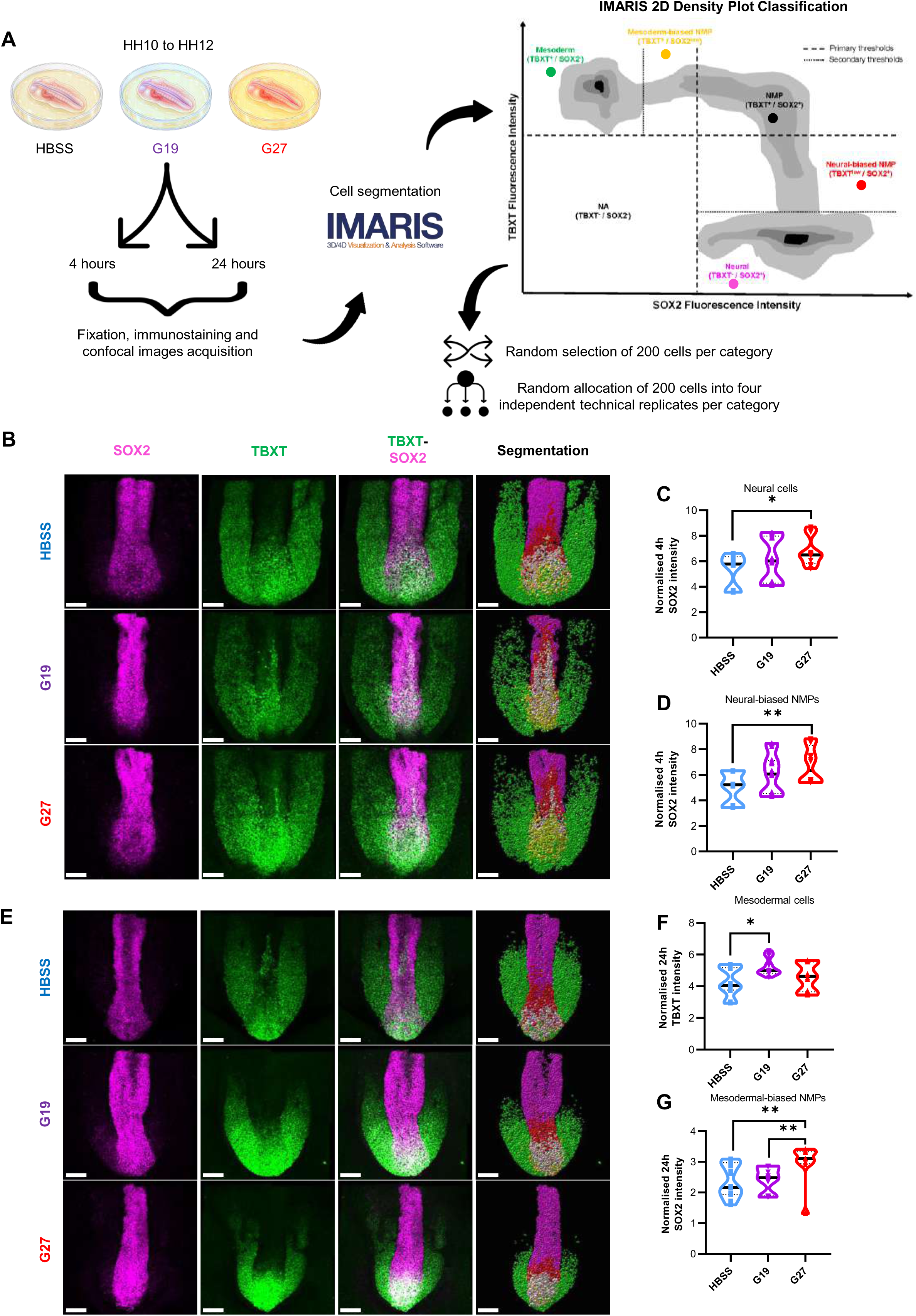
- Hemichannel and gap-junction inhibition differentially affect SOX2 and TBXT expression across the NMP continuum A, Experimental workflow for embryo treatment with HBSS, Gap19 (G19) or Gap27 (G27), followed by immunostaining, IMARIS-based cell segmentation and classification of cell subpopulations using two-dimensional density plots based on *TBXT* and *SOX2* fluorescence intensities. For each category, 200 cells were randomly selected and distributed across four technical replicates. **B–D,** IMARIS-processed images showing *SOX2*, *TBXT* and merged fluorescence signals, together with the corresponding cell subpopulation segmentation, at 4 h of treatment. Quantification of *SOX2* and *TBXT* fluorescence intensity across the identified subpopulations is shown. Statistical significance was assessed using the Kruskal–Wallis test followed by Dunn’s multiple-comparisons test with adjusted P values. At 4 h, the Kruskal–Wallis test indicated significant differences in *SOX2* intensity among neural cells (P = 0.0332) and neural-biased NMPs (P = 0.0123). Dunn’s multiple-comparisons test identified significant differences between HBSS and G27 in both neural cells (P = 0.0274) and neural-biased NMPs (P = 0.00910). n = 3, 5 and 4 embryos for HBSS, G19 and G27, respectively. **E–G,** IMARIS-processed images showing *SOX2*, *TBXT* and merged fluorescence signals, together with the corresponding cell subpopulation segmentation, at 24 h of treatment. Quantification of *SOX2* and *TBXT* fluorescence intensity across the identified subpopulations is shown. Statistical significance was assessed using the Kruskal–Wallis test followed by Dunn’s multiple-comparisons test with adjusted P values. At 24 h, the Kruskal–Wallis test indicated significant differences in *TBXT* intensity among mesodermal cells (P = 0.0235), with a significant difference between HBSS and G19 identified by Dunn’s multiple-comparisons test (P = 0.0215). Significant differences in *SOX2* intensity were also detected among mesodermal-biased NMPs (Kruskal–Wallis, P = 0.0050), with significant differences between G19 and G27 (P = 0.00980) and between HBSS and G27 (P = 0.00880). n = 5, 4 and 3 embryos for HBSS, G19 and G27, respectively.

After 4 h of treatment, SOX2 levels were significantly altered within specific progenitor populations. In neural cells, SOX2 expression was increased following GAP27 treatment relative to HBSS controls (Fig. 4B,C), and a similar effect was observed in neural-biased NMPs (Fig. 4B,D). By contrast, GAP19 treatment produced more modest changes and was not significantly different from HBSS in either population. These observations indicate that gap-junction inhibition rapidly shifts cells toward a stronger neural transcriptional state.

In contrast, TBXT expression was largely unaffected at this early time point. No significant differences were detected between either connexin inhibitor and HBSS controls in all the cell categories (supplementary figure 4). However, TBXT expression in neural cells differed significantly between GAP19- and GAP27-treated embryos, indicating that hemichannel and gap-junction inhibition produce distinct effects despite the absence of a detectable change relative to control conditions. Together, these results suggest that early responses to connexin inhibition primarily affect SOX2 expression while leaving TBXT levels largely unchanged.

We next examined whether these early transcriptional responses were maintained following 24 h of treatment. At this later stage, connexin inhibition produced distinct effects depending on both the cell population analysed and the mode of connexin inhibition.

The strongest effect on TBXT expression was observed in mesodermal cells, where GAP19 treatment significantly increased TBXT levels relative to HBSS controls (Fig. 4E,F). This effect was not observed following GAP27 treatment and represents the only significant difference between GAP19 and control embryos detected across the dataset. These results therefore suggest that prolonged inhibition of Cx43 hemichannels selectively affects mesodermal transcriptional program.

In contrast, SOX2 regulation was primarily associated with GAP27 treatment. Within the mesoderm-biased NMP population, SOX2 levels were significantly higher following GAP27 treatment than in either HBSS- or GAP19-treated embryos (Fig. 4E,G). Thus, while hemichannel inhibition promoted increased TBXT expression in mesodermal cells, gap-junction inhibition preferentially affected SOX2 expression within progenitor populations that retain both neural and mesodermal characteristics.

Interestingly, within the central NMP population, SOX2 levels differed significantly between GAP19 and GAP27 treatments despite neither treatment being significantly different from HBSS controls (supplementary Figure 4). Although this result does not support a detectable change relative to untreated embryos, it indicates that inhibition of hemichannels and gap junctions does not produce equivalent transcriptional responses within NMPs.

Taken together, these findings indicate that connexin inhibition does not induce a uniform transcriptional response across the NMP continuum. Instead, hemichannel and gap-junction inhibition affect distinct cell populations and different transcription factors. GAP19 primarily influences TBXT expression in mesodermal cells, whereas GAP27 preferentially alters SOX2 expression within progenitor populations. These observations suggest that the two modes of connexin-mediated communication contribute differently to the regulation of neural and mesodermal transcriptional programmes.

### Connexin inhibition disrupts posterior tissue organisation

To determine whether the molecular alterations induced by connexin inhibition were associated with changes in tissue organisation, we performed quantitative morphometric analyses of embryos following treatment with GAP19 and GAP27 (Fig. 5A–F). Measurements of global embryonic dimensions revealed only modest changes in total embryo length, posterior axis length, presomitic mesoderm size and neural tube width following connexin inhibition (Fig. 5A–D). Although trends toward reduced posterior growth were observed in treated embryos, these effects were variable between embryos and did not consistently reach statistical significance.

**Figure 5.**
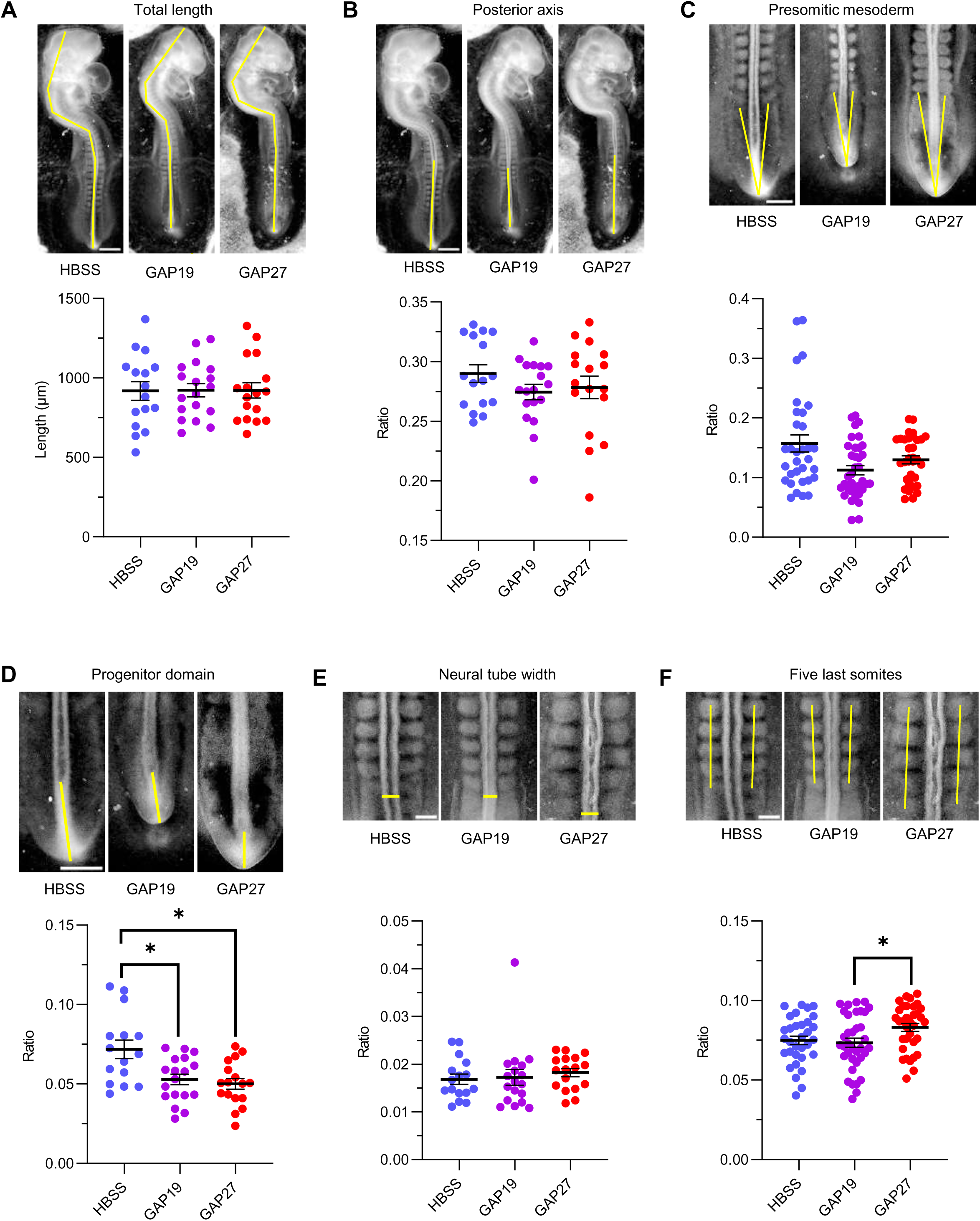
- Connexin inhibition disrupts posterior tissue organisation during axis elongation A, Representative images showing total embryonic length in embryos treated with HBSS, Gap19 (G19) or Gap27 (G27), with the measurement trajectory indicated by the yellow line. Quantification of total embryonic length is shown below, with individual data points displayed for each treatment condition. **B,** Representative images showing the measurement of the posterior embryonic axis in embryos treated with HBSS, Gap19 (G19) or Gap27 (G27), with the measurement trajectory indicated by the yellow line. Quantification of the posterior axis is shown below as the ratio of posterior axis length to total embryonic length. **C,** Representative images showing the measurement of the presomitic mesoderm (PSM) region in embryos treated with HBSS, Gap19 (G19) or Gap27 (G27), with measurement trajectories indicated by yellow lines. Quantification of the PSM region is shown below as the ratio of PSM length to total embryonic length. The Kruskal–Wallis test showed a marginal, but non-significant, difference among treatment groups (P = 0.0594). No significant pairwise differences were detected by Dunn’s multiple-comparisons test. **D,** Representative images showing the measurement of the progenitor domain in embryos treated with HBSS, Gap19 (G19) or Gap27 (G27), with the measurement trajectory indicated by the yellow line. Quantification of the progenitor domain is shown below as the ratio of progenitor domain length to total embryonic length. The Kruskal–Wallis test indicated a significant difference among treatment groups (P = 0.0120). Dunn’s multiple-comparisons test showed significant differences between HBSS and G19 (P = 0.0497) and between HBSS and G27 (P = 0.0166). **E,** Representative images showing the measurement of neural tube width in embryos treated with HBSS, Gap19 (G19) or Gap27 (G27), with the measurement indicated by the yellow line. Quantification of neural tube width is shown below as the ratio of neural tube width to total embryonic length. **F,** Representative images showing the measurement of the five most posterior somites in embryos treated with HBSS, Gap19 (G19) or Gap27 (G27), with measurement trajectories indicated by yellow lines. Quantification of the five most posterior somites is shown below as the ratio of the length of the five most posterior somites to total embryonic length. The Kruskal–Wallis test indicated a significant difference among treatment groups (P = 0.0367), with significant pairwise differences detected between G19 and G27 p=0,0495 after Dunn’s multiple-comparisons test. n = 16 (HBSS), n = 18 (G19) and n = 17 (G27); these sample sizes apply to all panels (A–F). Statistical significance was assessed using the Kruskal–Wallis test followed by Dunn’s multiple-comparisons test with adjusted P values.

In contrast, connexin inhibition produced clear alterations within the posterior growth zone. Quantification of the progenitor domain revealed a significant reduction in its relative size following connexin inhibition, indicating that connexin-mediated communication contributes to the maintenance of the neuromesodermal progenitor compartment during axis elongation (Fig. 5E). To determine whether these organisational defects extended to newly formed segmental structures, we analysed the length of the five most recently formed somites which were expected to be produced during the 24 hours treatment with the inhibitors. The length occupied by the five most recently formed somites differed significantly between treatment conditions, with GAP27-treated embryos displaying greater values than GAP19-treated embryos (Fig. 5F), demonstrating that perturbation of connexin-mediated cell t cell communication affects the spatial organisation of the posterior axis.

Together, these observations indicate that connexin activity contributes to the organisation of posterior tissues during axis elongation. While overall embryo morphology remains relatively preserved, disruption of connexin-mediated communication reduces the size of the progenitor compartment and alters newly formed somites, suggesting that communication-dependent mechanisms are required for maintaining posterior tissue architecture.

### Connexin activity is required for robust and coordinated somitogenesis

Given the defects in somite organisation and the central role of NMPs in sustaining somitogenesis, we next investigated whether connexin inhibition affects segmentation dynamics. Time-lapse imaging of control embryos revealed highly regular somite formation, with successive somites forming at predictable intervals (Fig. 6A,B). In contrast, embryos treated with GAP19 or GAP27 frequentlydisplayed abnormal segmentation events, characterised by prolonged pauses between somite formation or, conversely, by the rapid appearance of multiple somites within a short time interval (Fig. 6C–F).

**Figure 6.**
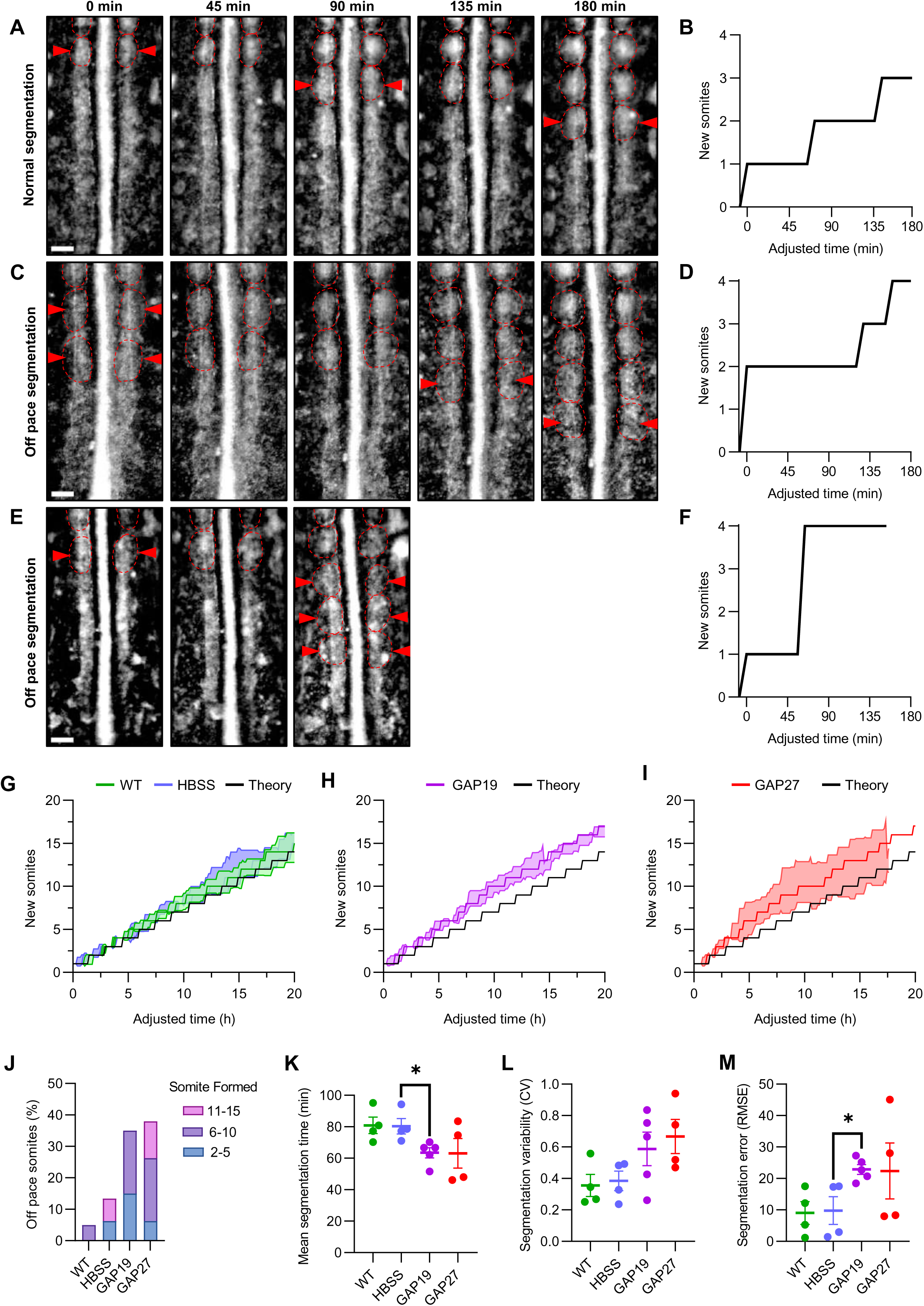
– Connexin activity is required for robust and coordinated somitogenesis A, **B**, Representative images and corresponding graph showing the cumulative number of newly formed somites (red arrowheads) as a function of adjusted time during normal somite segmentation. Dotted circles represent formed somites. **C, D,** Representative images and corresponding graph showing the cumulative number of newly formed somites (red arrowheads) as a function of adjusted time during delayed somite segmentation, in which somite formation occurs at a slower rate than under normal segmentation conditions. Dotted circles represent formed somites. **E, F,** Representative images and corresponding graph showing the cumulative number of newly formed somites (red arrowheads) as a function of adjusted time during altered somite segmentation, in which multiple somites are generated within the same time interval, resulting in a segmentation pattern that deviates from the normal timing of somite formation. Dotted circles represent formed somites. **G-I** Mean cumulative number of newly formed somites plotted as a function of adjusted time. Green, WT embryos; blue, HBSS-treated embryos; Purple, G19-treated embryos; Red, G27-treated embryos; black, theoretical expected somite formation curve. **J,** Percentage of embryos exhibiting off pace somite formation at the indicated somites categories (2– 5, 6–10 and 11–15 somites) in WT, HBSS, GAP19 and GAP27 embryos. **K,** Quantification of the average time required for somite formation in WT embryos and embryos treated with HBSS, GAP19 or GAP27. Individual data points correspond to independent embryos; bars represent mean ± s.e.m. Statistical significance was assessed using a two-tailed unpaired t-test indicating a significant difference between HBBS and G19 (P = 0.0206). **L,** Quantification of inter-embryo variability in segmentation timing for WT, HBSS-, GAP19- and GAP27-treated embryos. Individual data points correspond to independent embryos; bars represent mean ± s.e.m. **M,** Quantification of the root mean square error (RMSE) between observed and predicted segmentation dynamics showing segmentation error in WT-, HBSS-, GAP19-, GAP27-treated embryos. Statistical significance was assessed using a two-tailed unpaired t-test indicating a significant difference between HBBS and G19 (P = 0.0182).

To determine how these local segmentation defects affected the progression of somitogenesis, we next examined cumulative somite formation trajectories. While WT and HBSS embryos closely followed the expected theoretical segmentation programme, GAP19- and GAP27-treated embryos progressively diverged from this trajectory over time (Fig. 6G–I). This divergence was particularly apparent during later stages of segmentation, indicating a gradual loss of coordination in somitogenesis.

These qualitative observations were further reflected by an increase in the proportion of off-pace somites formed in connexin-inhibited embryos (Fig. 6J). Notably, abnormal segmentation events were already detected among the earliest somites analysed (somites 2–5), indicating that perturbation of connexin activity rapidly affects segmentation dynamics following treatment. However, the frequency of off-pace somites continued to increase during subsequent segmentation cycles and reached its highest level among the formation of somites 11–15. This progressive accumulation of abnormal events suggests that connexin activity is required continuously during ongoing somitogenesis rather than solely during establishment of the segmentation program. While GAP19 treatment induced an early increase in off-pace segmentation, GAP27-treated embryos tended to exhibit the highest frequency of abnormal events at later stages suggesting different role on somitogenesis regulation.

To quantify segmentation timing, we measured the mean segmentation time for each embryo. Segmentation dynamics differed significantly across experimental conditions (Fig. 6K). Post-hoc analyses revealed a significant reduction in segmentation time following GAP19 treatment relative to HBSS controls indicating that inhibition of Cx43 hemichannels accelerates somitogenesis. Although GAP27-treated embryos exhibited a similar trend, they did not significantly differ from HBSS controls. We next asked whether connexin inhibition affected the reproducibility of the segmentation program. Although the coefficient of variation (CV) of segmentation intervals tended to increase following connexin inhibition, this effect did not reach statistical significance (Fig. 6L), suggesting that altered segmentation dynamics cannot be explained solely by increased interval variability.

To directly assess segmentation robustness, we calculated the root mean squared error (RMSE) between observed and theoretical cumulative somite formation trajectories. RMSE values differed significantly across experimental conditions (Fig. 6M). Post-hoc comparisons identified a significant increase in RMSE following GAP19 treatment relative to HBSS controls, indicating that hemichannel inhibition leads to progressive deviations from the expected segmentation program. GAP27-treated embryos also displayed elevated RMSE values and greater dispersion of segmentation trajectories, although these differences did not reach statistical significance relative to HBSS controls.

Importantly, segmentation was never completely arrested, indicating that connexin activity is not required for somite formation itself but rather for maintaining the precision and reproducibility of the segmentation program. Together, these findings demonstrate that connexin-mediated communication contributes to both the timing and robustness of somitogenesis. Inhibition of Cx43 hemichannels is sufficient to significantly alter segmentation timing and increase deviation from the expected developmental program, whereas inhibition of gap-junction communication produces qualitatively similar but more variable effects. These observations suggest that the two modes of connexin-mediated communication contribute differently to the coordination of segmentation dynamics.

Taken together, these observations demonstrate that connexin-mediated communication contributes to both the temporal and spatial robustness of somitogenesis. While segmentation remains active following connexin inhibition, embryos progressively diverge from the expected developmental trajectory and display altered positioning of newly formed somite boundaries, resulting in a less coordinated and less reproducible segmentation program.

## DISCUSSION

In this study, we identify GJA1 as a core component of the neuromesodermal progenitor domain and demonstrate that connexin-43 (Cx43) is both enriched and functionally active in this population. The spatial and temporal pattern of GJA1 expression, together with the heterogeneous distribution of Cx43 protein and the increased permeability observed in the coumarin assay, indicate that neuromesodermal progenitors constitute a highly communication-competent cell population. While the regulation of NMP states has primarily been studied through transcriptional networks and signalling pathways such as WNT, FGF and retinoic acid, our findings suggest that intercellular communication represents an additional and previously underappreciated layer of regulation contributing to the organisation and behaviour of the NMP continuum within the posterior axis (Goodenough & Paul, 2003; Evans & Martin, 2002; Solan & Lampe, 2009; Laird, 2014).

A central aspect of this work is the distinction between the two modes of connexin-43 activity: gap junctions, which mediate direct cell–cell coupling, and hemichannels, which allow exchange between the intracellular and extracellular environment. Using pharmacological inhibitors with distinct specificities, we found that these two modes of communication affect different cellular populations and transcriptional programs. Gap-junction inhibition primarily altered SOX2 expression within neural-biased and mesoderm-biased NMP populations, whereas hemichannel inhibition produced the strongest effect on TBXT expression in mesodermal cells following prolonged treatment. These observations indicate that hemichannels and gap junctions do not contribute equivalently to the regulation of cell states across the NMP continuum and instead appear to influence distinct transcriptional programs.

The coumarin assay further supports this interpretation. The rapid loss of tracer from the NMC domain in the absence of detectable accumulation within neighbouring cells is consistent with a substantial contribution of connexin-mediated exchange with the extracellular environment. Together, these findings suggest that hemichannel activity may contribute to shaping the local signalling environment of the posterior growth zone, whereas gap junctions may play a more prominent role in coordinating responses between neighbouring cells, consistent with the distinct signalling functions described for connexin hemichannels and gap junctions in other developmental and physiological contexts (Wang et al., 2013; Abudara et al., 2014; Laird, 2014).

One of the most striking findings of this study is that inhibition of connexin activity produced distinct effects on SOX2 and TBXT expression depending on treatment duration and cell population. Early responses were dominated by alterations in SOX2 expression, whereas TBXT remained largely unchanged. At later stages, however, prolonged hemichannel inhibition induced increased TBXT expression in mesodermal cells, while gap-junction inhibition preferentially affected SOX2 expression within mesoderm-biased NMP populations.

These observations suggest that neural and mesodermal transcriptional programmes can be perturbed independently, at least transiently. Such behaviour differs from simplified models in which SOX2 and TBXT act as strictly antagonistic regulators and instead supports the view that NMPs exist as a continuum of transcriptional states shaped by multiple regulatory inputs (Tsakiridis and Wilson, 2015; Gouti et al., 2017; Koch et al., 2017; Oginuma et al., 2017; Verrier et al., 2018; Dias-Cuadros et al., 2020; Wymeersch et al., 2021). Connexin-mediated communication may therefore represent an additional regulatory layer capable of influencing specific components of this network without producing an immediate reciprocal response in the corresponding lineage marker.

At the morphogenetic level, connexin inhibition alters the organisation of the posterior growth zone. The most consistent defects are observed within the neuromesodermal progenitor compartment, whose relative size is significantly reduced following connexin inhibition. In addition, newly formed somites were also affected and longer in the GAP-27 inhibited embryos, indicating that connexin-mediated communication contributes to the spatial organisation of posterior tissues. Together, these observations suggest that connexin activity supports the architecture of the posterior growth zone during axis elongation. The phenotypes observed following hemichannel inhibition are consistent with the idea that exchange with the extracellular environment contributes to maintaining the organisation of posterior progenitor populations.

The most striking phenotype observed upon connexin inhibition concerns the disruption of segmentation dynamics. Control embryos display highly regular somite formation and closely follow the expected segmentation program, whereas connexin-inhibited embryos exhibit irregular segmentation characterised by prolonged pauses, accelerated segmentation events and progressive divergence from the theoretical trajectory. Importantly, off-pace somites appear rapidly following connexin inhibition and accumulate during subsequent segmentation cycles, indicating that connexin-mediated communication contributes continuously to segmentation dynamics rather than solely during the establishment of the segmentation program. This behaviour differs from classical modulation of segmentation clock speed, which typically produces uniform acceleration or deceleration while preserving periodicity (Hubaud et al., 2017). Instead, the defects observed here reflect a reduction in developmental robustness, defined as the ability of the segmentation system to maintain reproducible patterning despite intrinsic developmental noise, and suggest a breakdown in the coordination of oscillatory activity at the tissue level (Oates et al., 2012; Hubaud et al., 2017; Sonnen et al., 2018).

Consistent with these temporal defects, connexin inhibition also alters the spatial organisation of segmentation. In particular, the five most recently formed somites occupy a greater axial length following GAP27 treatment, indicating that disruption of connexin-mediated communication affects the organisation of newly generated segments. Because these somites are formed during the treatment period, this observation suggests that connexin activity contributes to the spatial organisation of ongoing segmentation. Within the classical framework of somitogenesis, segment geometry emerges from the coupling between segmentation dynamics and tissue growth (Dubrulle & Pourquié, 2004; Gomez et al., 2008; Lauschke et al., 2013; Meijer and Sonnen, 2024). The altered somite organisation observed here therefore likely reflects reduced coordination between these processes, reinforcing the idea that connexin-mediated communication contributes not only to temporal precision but also to the spatial robustness of segmentation.

Such irregular segmentation and altered somite geometry have previously been associated with defects in intercellular coupling. In particular, Notch signalling plays a central role in synchronising oscillations between neighbouring presomitic mesoderm cells, and disruption of this coupling leads to desynchronised segmentation dynamics and irregular somite formation (Jiang et al., 2000; Dequeant and Pourquié, 2008; Lewis et al., 2009; Oates et al., 2012). Our results extend this framework by identifying connexin-43-mediated communication as an additional mechanism contributing to oscillatory coupling. The rapid appearance and progressive accumulation of off-pace segmentation events following connexin inhibition suggest that oscillatory precision depends on continuous intercellular communication throughout active somitogenesis.

To our knowledge, connexin-mediated communication has not previously been directly implicated in the control of somitogenesis or segmentation clock dynamics. Intercellular synchronisation in the presomitic mesoderm has been largely attributed to Notch signalling, yet whether additional mechanisms contribute to this coordination has remained unclear. Our results therefore identify connexin-43 as a previously unrecognised regulator of segmentation clock robustness, and suggest that multiple layers of intercellular communication act in concert to ensure coherent oscillatory behaviour during axis elongation (Oates et al., 2012; Sonnen et al., 2018; Meijer and Sonnen, 2024).

In this framework, gap junctions are well suited to support direct synchronisation between adjacent cells (Evans & Martin, 2002; Goodenough & Paul, 2003; Wang et al., 2013; Isomura et al., 2026), whereas hemichannels may regulate extracellular signals, such as ions or metabolites, that influence oscillatory dynamics at a broader scale. Together, disruption of either communication mode reduces temporal and spatial robustness observed in connexin-inhibited embryos.

Taken together, our findings support a model in which connexin-43 links regulation of neuromesodermal progenitor states to the robustness of segmentation during vertebrate axis elongation. Within the posterior growth zone, connexin-mediated communication contributes to the organisation of the NMP continuum and influences distinct transcriptional programs through hemichannel- and gap-junction-dependent mechanisms, supporting the growing view that connexins act not only as channels for intercellular exchange but also as regulators of developmental signalling and tissue organisation (Laird, 2014). At the tissue level, this communication supports the organisation of posterior structures and maintains the reproducibility of segmentation dynamics. The disruption of connexin activity therefore affects multiple levels of organisation simultaneously, linking alterations in progenitor states to defects in developmental timing and spatial patterning. These findings suggest that intercellular communication does not simply modulate individual developmental processes, but instead contributes to the coordination of multiple levels of organisation, linking progenitor-state regulation to the temporal and spatial robustness of embryonic pattern formation.

## MATERIALS AND METHODS

### Ethics statement

Fertilized White Leghorn chicken eggs were obtained from a local supplier and used exclusively for embryological studies prior to hatching. Experimental procedures were conducted in accordance with institutional guidelines and applicable French and European regulations concerning the use of avian embryos in research. No hatched animals were used in this study.

### Embryo collection and staging

Fertilized White Leghorn chicken eggs were obtained from a local biological farm (EARL Les Bruyères, Centre Val de Loire) and incubated at 38°C in a humidified incubator (Borotto, N.1 Real 49) maintained at 60–70% relative humidity. Embryos were harvested by opening the eggs and transferring them onto Whatman filter paper using the filter paper carrier technique described by Chapman et al. (2001). Embryonic developmental stages were determined according to the criteria established by Hamburger and Hamilton (1992).

### Embryo culture and pharmacological treatments

For *ex ovo* culture experiments, agarose–albumen culture plates were prepared by dissolving 0.3 g Bacto Agar (Sigma-Aldrich, A5306) and 0.8 mL of 20% D-Glucose (Sigma-Aldrich, G8270) in a mixture containing 25 mL egg albumen and 1.23 mL 5M NaCl, brought to a final volume of 50 mL with pasteurized osmotic water. Approximately 2 mL of the agarose–albumen mixture was dispensed into 35 mm culture dishes (Falcon, 353001) and stored at 4°C until use.

Treatment solutions were prepared using Hanks’ Balanced Salt Solution (HBSS, Thermo Fisher Scientific, 14025092) as vehicle control. GAP19 (Sigma-Aldrich, 1507930-57-5) and GAP27 (Sigma-Aldrich, 198284-64-9) were diluted at 3:1000 in HBSS (final concentration of 3μg/mL). For each experiment, 50 µL of treatment solution was applied to the culture substrate prior to embryo transfer, followed by an additional 50 µL applied directly onto the embryo to ensure continuous exposure.

Embryos were cultured at 38°C in a humid chamber for the durations specified in each experiment before fixation or live imaging.

### Whole-mount HCR-Fluorescence In Situ Hybridization (HCR-FISH)

Whole-mount RNA Hybridization Chain Reaction Fluorescence In Situ Hybridization (HCR-FISH) was performed according to the manufacturer’s instructions (Molecular Instruments) and previously published protocols (Choi et al., 2018).

Embryos were fixed in 4% paraformaldehyde (PFA; Electron Microscopy Sciences) for 1 h at room temperature, dehydrated and stored in 100% methanol at −20°C. Samples were subsequently rehydrated in graded methanol/PBST solutions (75/25, 50/50, 25/75, 0/100) and permeabilized with Proteinase K (10µg/ml) (Sigma-Aldrich, P6556). Following a post-fixation step in 4% PFA, embryos were equilibrated in 5× SSCT and pre-hybridized in hybridization buffer for 30 min at 37°C. Hybridization was carried out overnight (12–14 h) at 37°C using probe sets directed against GJA1, SOX2 and TBXT (SOX2 Accession # NM_205188.3; TBXT Accession # NM_204940.2; GJA1 Accession # NM_204586.3). Following hybridization, embryos were washed in probe wash buffer and 5× SSCT according to the manufacturer’s instructions. Samples were then pre-amplified and incubated overnight with snap-cooled fluorescent hairpins.

Following amplification, embryos were washed extensively in 5× SSCT, post-fixed in 4% PFA and stored in PBS until imaging.

Images were acquired on a Zeiss LSM980 laser-scanning confocal microscope using a 10X or 40× objective.

### Quantification of HCR signal

Fluorescence intensity profiles were quantified using Fiji (ImageJ) using the 10X images. For each embryo, a line region of interest (ROI) spanning the neural plate, NMC domain and primitive streak was manually defined. Fluorescence intensity values were extracted using the Plot Profile function and normalized to the maximal intensity measured within each embryo.

### Immunofluorescence staining

Embryos were fixed overnight at 4°C in 4% PFA and dissected in PBS under a stereomicroscope. Samples were washed three times in PBST (PBS supplemented with 0.1% Triton X-100) and blocked for a minimum of 1 h in PBST containing 10% fetal bovine serum (FBS).

Embryos were incubated overnight at 4°C with primary antibodies diluted in blocking solution. The <u>following antibodies were used:</u>

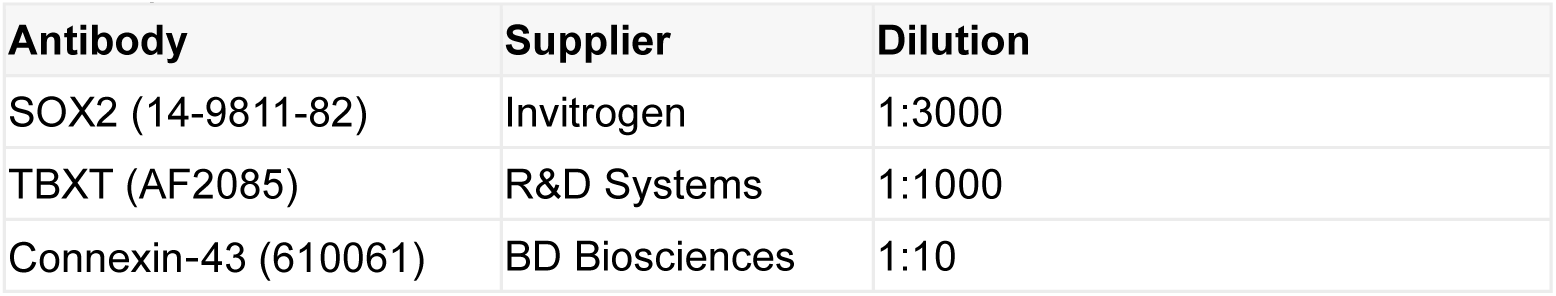

Following three washes in PBST, embryos were incubated overnight with appropriate fluorophore-conjugated secondary antibodies.

### Per-cell Cx43 puncta detection and analysis

High-resolution immunofluorescence images were acquired using a Zeiss LSM980 confocal microscope equipped with a 63× objective. In the dorsal region of each embryo, square regions of interest (134.7 × 134.7 µm) were selected within neural, mesodermal, and neuromesodermal progenitor (NMP) tissues based on tissue morphology and immunoreactivity for TBXT and SOX2. Image stacks consisted of more than 20 optical sections acquired at 0.4 µm intervals along the z-axis.

To increase the number of cells available for analysis while minimizing signal overlap, two spatially separated cellular layers were selected from each z-stack. For each layer, a mean-intensity projection was generated from three consecutive z-planes, resulting in one dorsal and one ventral projected image per stack.

Because phalloidin staining produced a discontinuous and noisy labeling of cell boundaries, cell identification was based on the nuclear markers TBXT and SOX2. Nuclei were segmented using CellProfiler and subsequently expanded using a Voronoi-like propagation approach to approximate cellular territories. To avoid overexpansion artifacts, only cells whose propagated area increased less than 3.5-fold relative to the original nuclear area were retained for downstream analyses.

Cx43 puncta were identified using a single Ilastik pixel-classification project trained and applied uniformly across all images to ensure consistent object detection. To reduce false-positive assignments, only puncta larger than five pixels were considered for further analysis.

Detected puncta were subsequently dilated by six pixels and overlaid onto the Voronoi-expanded cellular territories. Each punctum was assigned to the corresponding CellID according to spatial overlap. Puncta associated with a single CellID were classified as putative Cx43 hemichannels, whereas puncta overlapping two neighboring CellIDs were classified as intercellular Cx43 channels (gap junctions). Quantitative measurements were then extracted on a per-cell basis for subsequent statistical analyses.

### Photoactivatable coumarin permeability assay

Posterior embryonic regions were loaded with a photoactivatable coumarin-based tracer and locally photoactivated using defined regions of interest. Time-lapse acquisition was conducted on a Zeiss LSM980 confocal microscope under constant imaging conditions.

Fluorescence intensity was measured over time within the activated region. Signal loss was quantified and normalized to the initial fluorescence intensity (F/F0). Rates of fluorescence decay were compared between neural, NMC and mesodermal regions.

### Analysis of NMP states following connexin inhibition

Embryos were cultured in WT, HBSS, GAP19 or GAP27 conditions for either 4 h or 24 h before fixation and immunostaining.

Single-cell segmentation was performed using Imaris (Oxford Instruments). TBXT and SOX2 fluorescence channels were preprocessed using a rolling-ball background subtraction (radius= 50 µm) followed by Gaussian filtering (σ=0.389 µm, corresponding to one voxel). The processed TBXT and SOX2 channels were then combined using a logical AND operation, and used to generate a segmentation mask. SOX2 and TBXT fluorescence intensities were subsequently extracted from the original, unprocessed images for each segmented cell and visualized as a two-dimensional density plot representing SOX2 fluorescence intensity (x-axis) as a function of TBXT fluorescence intensity (y-axis).

The resulting density plots were analysed independently for each embryo. Primary thresholds corresponding to the lower boundaries of the SOX2 high expressing cells and TBXT high expressing cells populations were used to partition the plots into four quadrants: SOX2-/TBXT-−, SOX2+/TBXT-, SOX2+/TBXT-, and SOX2+/TBXT+.Cells located within the SOX2+/TBXT+ quadrant was classified as the “NeuroMesodermal Progenitor“ (NMP), whereas cells within the SOX2-/TBXT-quadrant were excluded from further analysis (NA).

To further resolve progenitor states, the SOX2-/TBXT+ and SOX2+/TBXT-quadrants were subdivided using secondary thresholds determined from the boundaries of the corresponding expression clusters within each embryo. Within the SOX2-/TBXT+ region, cells exhibiting high TBXT and low SOX2 expression were classified as mesodermal cells (TBXT⁺/SOX2⁻), whereas cells occupying the intermediate region between mesodermal cells and the NMP population were classified as mesoderm-biased NMPs (TBXT⁺/SOX2^low^)

Similarly, within the SOX2+/TBXT-region, cells exhibiting high SOX2 and low TBXT expression were classified as neural cells (SOX2⁺/TBXT⁻), whereas cells occupying the intermediate region between neural cells and the NMP population were classified as neural-biased NMPs (SOX2⁺/TBXT^low^).

The same hierarchical classification strategy was applied independently to each embryo, allowing the identification of neural cells, neural-biased NMPs, NMPs, mesoderm-biased NMPs and mesodermal cells while accounting for embryo-to-embryo variability in staining intensity. Cell population boundaries were manually positioned based on density-cluster distributions and visually validated for each embryo prior to downstream analysis.

Following cell segmentation and classification into the different categories, 200 cells were randomly selected from each category using a randomization software. The selected cells were subsequently randomly allocated into four independent subsets of 50 cells each, with each subset considered a technical replicate for subsequent statistical analyses.

For each cell population, SOX2 and TBXT fluorescence intensities were quantified and compared across treatment conditions.

### Morphometric analyses

Embryos collected at HH7^-^ to HH8, HH10 to HH12 and HH18 to HH20 of development were imaged using a TOMLOV TriL110 macroscope. All measurements were performed using identical anatomical landmarks across embryos such as forebrain, midbrain, hindbrain, neural tube, primitive streak, somite, tailbud, vitelline artery, Hensen’s Node, posterior limb buds

Measurements were performed in Fiji and included:

- Total embryo length: following the neural tube curvature: HH7^-^ to HH8 forebrain to end of primitive streak, HH10 to HH12/ HH18 to HH20 midbrain to tailbud
- Posterior axis length HH7^-^ to HH8 Hensen’s Node to tailbud, HH10 to HH12/ HH18 to HH20 vitelline artery to tailbud
- NMC domain length: HH10 to HH12/ HH18 to HH20: Hensen’s Node to tailbud
- Presomitic mesoderm length all: last somite to end of primitive streak/tailbud
- Neural tube width HH7^-^ to HH8/ HH10 to HH12 under last somite formed, HH18 to HH20 under posterior limb buds
- Tail width HH7^-^ to HH8/ HH10 to HH12 under last somite formed, HH18 to HH20 under posterior limb buds

### Heartbeat analysis

Embryos cultured under WT, HBSS, GAP19 and GAP27 conditions were imaged using a Leica THUNDER Model Organism imaging system (total movie duration: 2min; interval: 0.035s; total images: 3429; exposition: 10ms; binning: 2x2; objective: 1X/0,07; zoom: 5,09x and total magnification: 5,1x). Cardiac activity was recorded immediately following a 30 min stabilization period after addition of the pharmacological inhibition and again after 6 h of treatment.

Kymographs were generated from line ROIs crossing the cardiac tissue. Heartbeat periods were quantified using custom R scripts.

### Live imaging of segmentation dynamics

Embryos were cultured on agarose–albumen plates in the presence of WT, HBSS, GAP19 or GAP27 treatments.

After a stabilization period of 30 min, embryos were imaged every 8 min over a 24 h period using a Leica THUNDER Model Organism imaging system (total movie duration: 24h; interval: 8,012min; total images: 181; exposition: 50ms; binning: 2x2; objective: 1X/0,08; zoom: 6,38x and total magnification: 4,1x; tiles: from 2x6 to 3x6). Time-lapse sequences were tiled and exported as 8-bit AVI files for analysis.

### Quantification of somite formation

Somite formation events were manually annotated using the appearance of complete epithelial boundaries as reference points.

For each embryo, segmentation period was calculated as:

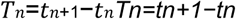

Where,

#### tntn

corresponds to the time of formation of somite

The following parameters were extracted:

- Mean segmentation period calculated as the average interval between the formation of successive somites across the first fifteen segmentation events recorded for each embryo.
- Standard deviation of segmentation intervals calculated from the distribution of the fifteen successive segmentation intervals for each embryo.
- Coefficient of variation (CV) calculated as the ratio between the standard deviation and the mean segmentation period (CV = SD/mean).
- Percentage of off-pace somites calculated as the proportion of somites forming outside the expected segmentation window. The expected segmentation period was defined from the average of untreated embryos and embryos treated with HBSS, for which the mean segmentation interval was 78 min with a standard deviation of 19.5 min. Somites were therefore classified as off-pace when the interval between two successive segmentation events was shorter than 39 min (mean − 2 SD) or longer than 117 min (mean + 2 SD). The percentage of off-pace somites was calculated for each embryo as:

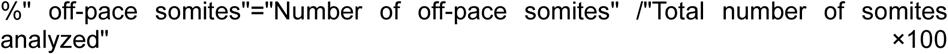

- Cumulative somite number over time obtained by plotting the total number of segmented somites as a function of time for each embryo.

#### Somite morphometry

Somite size and intersomite spacing were measured from time-lapse images and endpoint images using Fiji. Measurements were used to evaluate the spatial robustness of segmentation under connexin inhibition.

## Statistical analyses

All statistical analyses were performed using GraphPad Prism (version 10.5.0) and R (version 2024.09.0).

Data distributions were assessed prior to statistical testing. Comparisons involving more than two groups were analyzed using Kruskal–Wallis tests followed by Dunn’s multiple-comparison post hoc tests. Statistical significance was defined as:

- ns: p > 0.05
- *: p ≤ 0.05
- **: p ≤ 0.01
- ***: p ≤ 0.001
- ****: p ≤ 0.0001

Unless otherwise stated, each biological replicate corresponds to an independent embryo. Data are presented as mean ± SEM.

## Acknowledgements

The authors thank the scientists at the Institute of Genetics and Reproduction for insightful discussions, valuable suggestions, and constructive comments. We are particularly grateful to the Chazaud team for their contribution and support throughout this project. We also acknowledge the outstanding technical support provided by the Institute of Genetics, Reproduction and Development core facilities, including the Bioinformatics Platform (BIM) and the Clermont Confocal Imaging Platform (CLIC). Finally, we thank the institute’s administrative and technical support staff for their continued assistance.

---

**Supp Data Fig. 2.**
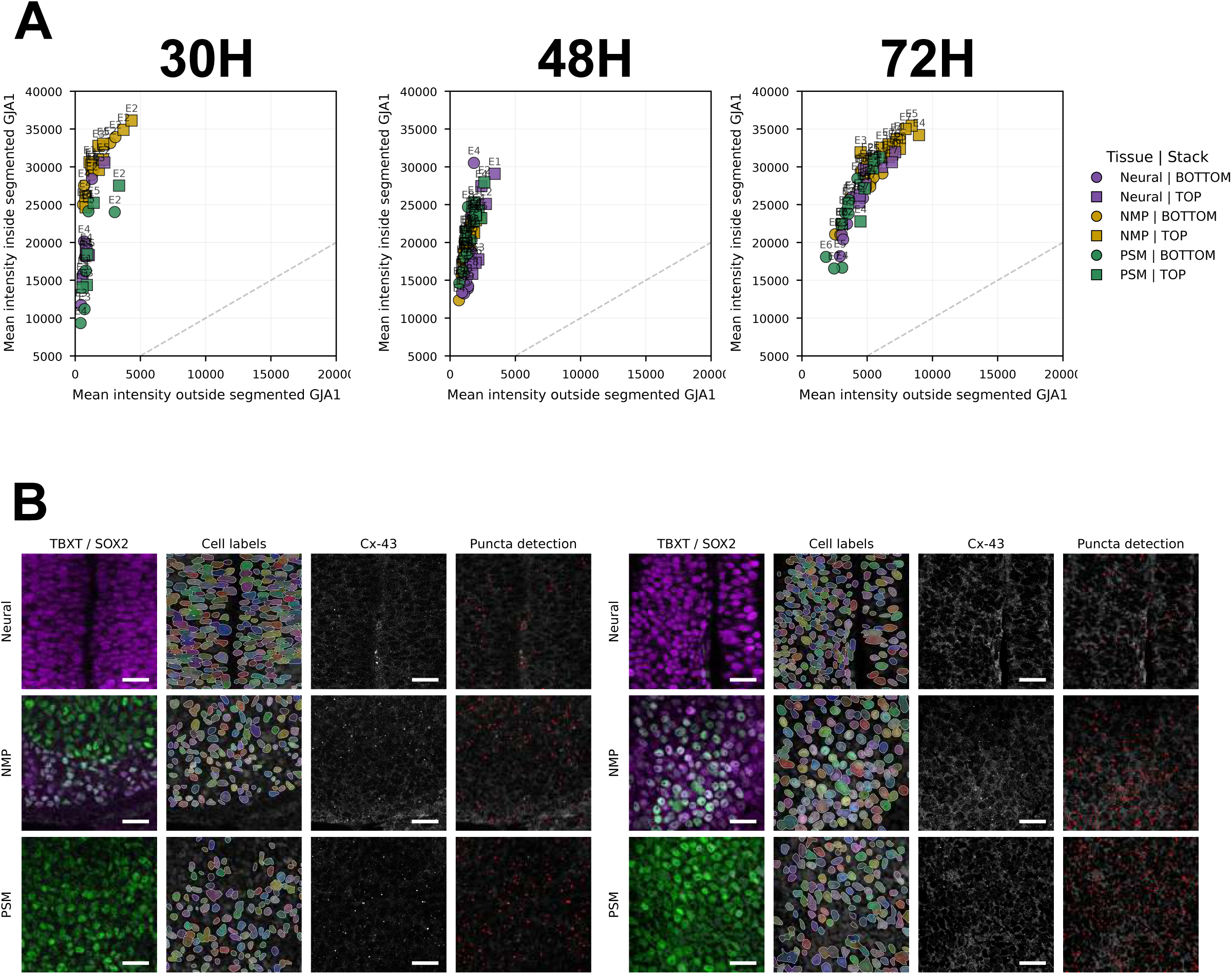
Cx43 puncta assignation in neural, mesodermal and NMP domains during body axis elongation. A) mean of GJA1 intensity in the segmentation mask obtained by ilastik versus mean intensity outside of the segmented mask. B) representative images of cell label assignation by cellprofiler and puncta detection by ilastik for stage HH10-12 (left) and HH18-20 (right)

**Supp Data Fig. 3.**
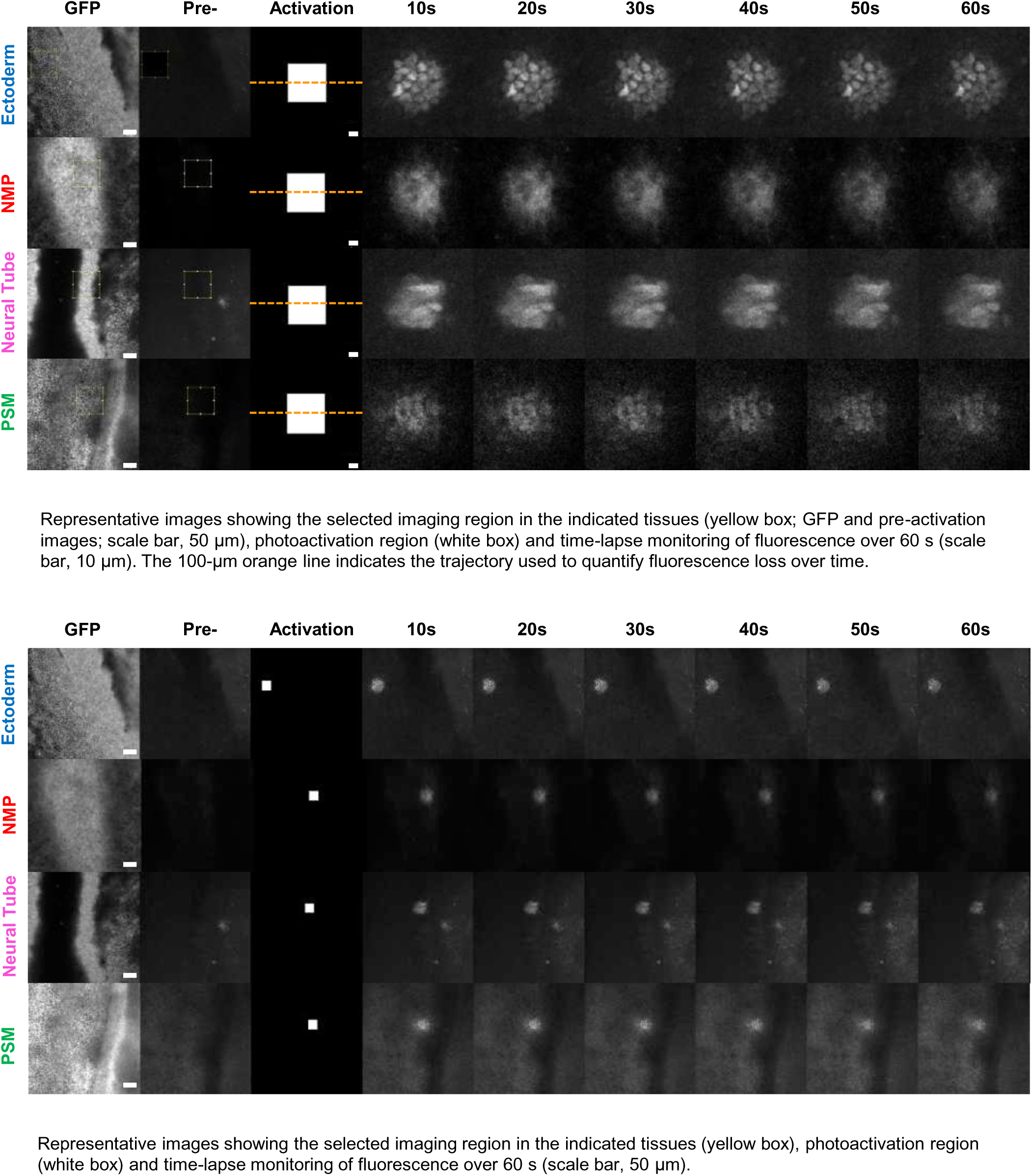
Representative images showing the selected imaging region in the indicated tissues (yellow box; GFP and pre-activation images; scale bar, 50 μm), photoactivation region (white box) and time-lapse monitoring of fluorescence over 60 s (scale bar, 10 μm). The 100-μm orange line indicates the trajectory used to quantify fluorescence loss over time. Representative images showing the selected imaging region in the indicated tissues (yellow box), photoactivation region (white box) and time-lapse monitoring of fluorescence over 60 s (scale bar, 50 μm).

**Supp Data Fig. 4.**
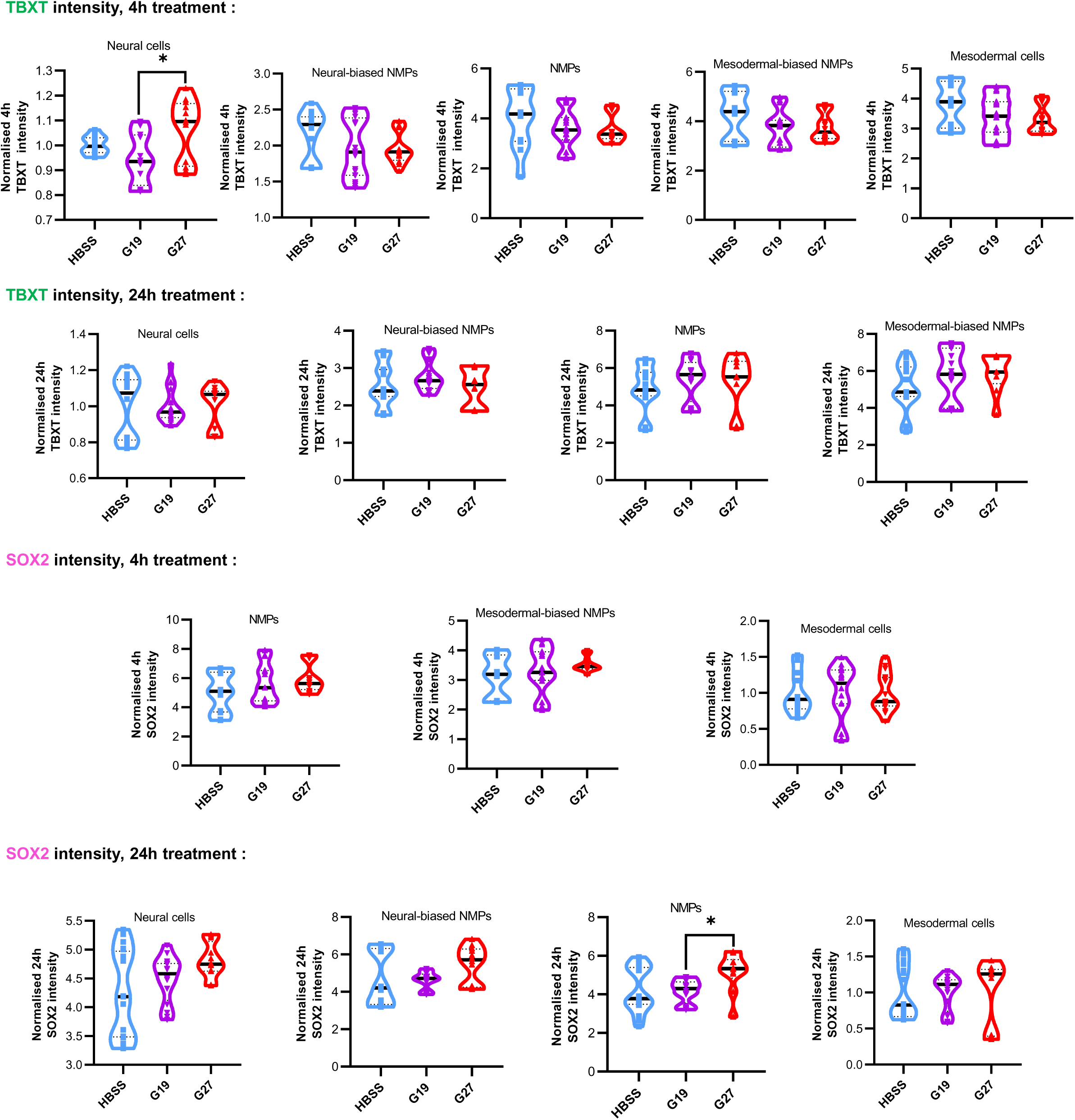
Quantification of SOX2 and TBXT fluorescence intensity across the identified subpopulations is shown (neural cells ; neural-biased NMPs ; NMPs ; mesodermal-biased NMPs ; mesodermal cells). Statistical significance was assessed using the Kruskal– Wallis test followed by Dunn’s multiple-comparisons test. At 4h, the Kruskal–Wallis test indicated significant differences in TBXT intensity among neural cells (P = 0,0484), with a significant difference between G19 and G27 identified by Dunn’s multiple-comparisons test (P = 0.0416). At 24 h, the Kruskal–Wallis test indicated significant differences in SOX2 intensity among NMPs (P = 0.0305), with a significant difference between G19 and G27 identified by Dunn’s multiple-comparisons test (P = 0.0334).

**Supp Data Fig. 6.**
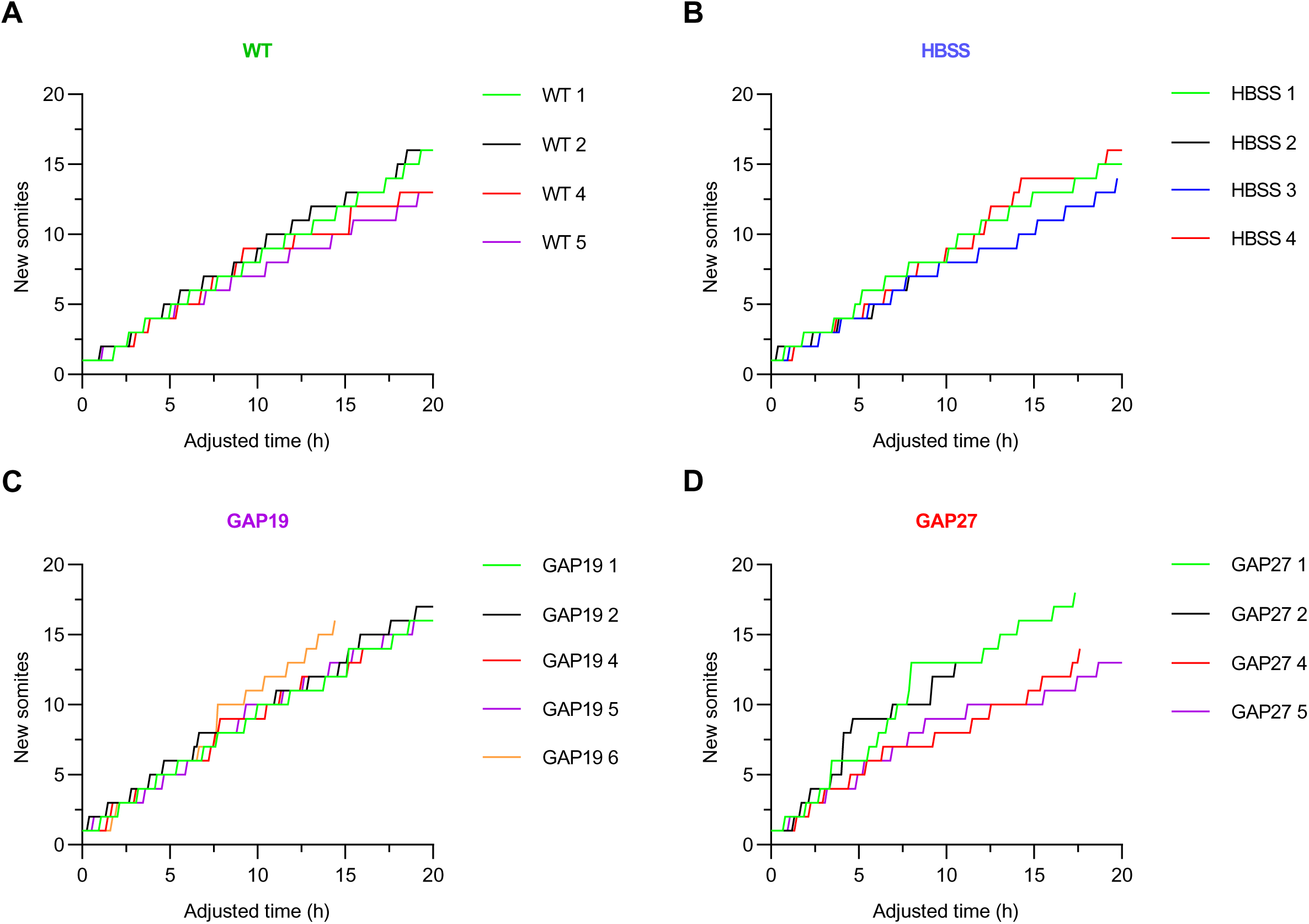
Mean cumulative number of newly formed somites plotted as a function of adjusted time per embryo after different treatments. A. No treatment, B. HBSS (control), C. GAP19 and D. GAP27.

## References

1. Abudara, V., Bechberger, J., Freitas-Andrade, M., De Bock, M., Wang, N., Bultynck, G., Naus, C. C., Leybaert, L., & Giaume, C. (2014). The connexin43 mimetic peptide Gap19 inhibits hemichannels without altering gap junctional communication in astrocytes. Frontiers in cellular neuroscience, 8, 306. 10.3389/fncel.2014.00306

2. Attardi, A., Fulton, T., Florescu, M., Shah, G., Muresan, L., Lenz, M. O., Lancaster, C., Huisken, J., van Oudenaarden, A., & Steventon, B. (2018). Neuromesodermal progenitors are a conserved source of spinal cord with divergent growth dynamics. *Development (Cambridge*, England), 145(21), dev166728. 10.1242/dev.166728

3. Binagui-Casas, A., Dias, A., Guillot, C., Metzis, V., & Saunders, D. (2021). Building consensus in neuromesodermal research: Current advances and future biomedical perspectives. Current opinion in cell biology, 73, 133–140. 10.1016/j.ceb.2021.08.003

4. Chapman, S. C., Collignon, J., Schoenwolf, G. C., & Lumsden, A. (2001). Improved method for chick whole-embryo culture using a filter paper carrier. Developmental dynamics : an official publication of the American Association of Anatomists, 220(3), 284–289. 10.1002/1097-0177(20010301)220:3<284::AID-DVDY1102>3.0.CO;2-5

5. Dequéant, M. L., & Pourquié, O. (2008). Segmental patterning of the vertebrate embryonic axis. Nature reviews. Genetics, 9(5), 370–382. 10.1038/nrg2320

6. Dubrulle, J., & Pourquié, O. (2004). Coupling segmentation to axis formation. *Development (Cambridge*, England), 131(23), 5783–5793. 10.1242/dev.01519

7. Diaz-Cuadros, M., Wagner, D. E., Budjan, C., Hubaud, A., Tarazona, O. A., Donelly, S., Michaut, A., Al Tanoury, Z., Yoshioka-Kobayashi, K., Niino, Y., Kageyama, R., Miyawaki, A., Touboul, J., & Pourquié, O. (2020). In vitro characterization of the human segmentation clock. Nature, 580(7801), 113–118. 10.1038/s41586-019-1885-9

8. Diaz-Cuadros, M., Miettinen, T.P., Skinner, O.S. et al. (2023). Metabolic regulation of species-specific developmental rates. Nature 613, 550–557. 10.1038/s41586-022-05574-4

9. Evans, W. H., & Boitano, S. (2001). Connexin mimetic peptides: specific inhibitors of gap-junctional intercellular communication. Biochemical Society transactions, 29(Pt 4), 606–612. 10.1042/bst0290606

10. Evans, W. H., & Martin, P. E. (2002). Gap junctions: structure and function (Review). Molecular membrane biology, 19(2), 121–136. 10.1080/09687680210139839

11. Ewart, J. L., Cohen, M. F., Meyer, R. A., Huang, G. Y., Wessels, A., Lo, C. W., & Fishman, M. C. (1997). Heart and neural tube defects in transgenic mice overexpressing the Cx43 gap junction gene. Development, 124(7), 1281–1292. 10.1242/dev.124.7.1281

12. Frith, T. J., Granata, I., Wind, M., Stout, E., Thompson, O., Neumann, K., Stavish, D., Heath, P. R., Ortmann, D., Hackland, J. O., Anastassiadis, K., Gouti, M., Briscoe, J., Wilson, V., Johnson, S. L., Placzek, M., Guarracino, M. R., Andrews, P. W., & Tsakiridis, A. (2018). Human axial progenitors generate trunk neural crest cells in vitro. eLife, 7, e35786. 10.7554/eLife.35786

13. Gomez, C., Ozbudak, E. M., Wunderlich, J., Baumann, D., Lewis, J., & Pourquié, O. (2008). Control of segment number in vertebrate embryos. Nature, 454(7202), 335–339. 10.1038/nature07020

14. Goodenough, D. A., & Paul, D. L. (2003). Beyond the gap: functions of unpaired connexon channels. Nature reviews. Molecular cell biology, 4(4), 285–294. 10.1038/nrm1072

15. Gouti, M., Tsakiridis, A., Wymeersch, F. J., Huang, Y., Kleinjung, J., Wilson, V., & Briscoe, J. (2014). In vitro generation of neuromesodermal progenitors reveals distinct roles for wnt signalling in the specification of spinal cord and paraxial mesoderm identity. PLoS biology, 12(8), e1001937. 10.1371/journal.pbio.1001937

16. Gouti, M., Delile, J., Stamataki, D., Wymeersch, F. J., Huang, Y., Kleinjung, J., Wilson, V., & Briscoe, J. (2017). A Gene Regulatory Network Balances Neural and Mesoderm Specification during Vertebrate Trunk Development. Developmental cell, 41(3), 243–261.e7. 10.1016/j.devcel.2017.04.002

17. Guillot, C., Djeffal, Y., Michaut, A., Rabe, B., & Pourquié, O. (2021). Dynamics of primitive streak regression controls the fate of neuromesodermal progenitors in the chicken embryo. eLife, 10, e64819. 10.7554/eLife.64819

18. Hamburger, V., & Hamilton, H. L. (1992). A series of normal stages in the development of the chick embryo. 1951. Developmental dynamics : an official publication of the American Association of Anatomists, 1S5(4), 231–272. 10.1002/aja.1001950404

19. Henrique, D., Abranches, E., Verrier, L., & Storey, K. G. (2015). Neuromesodermal progenitors and the making of the spinal cord. *Development (Cambridge*, England), 142(17), 2864–2875. 10.1242/dev.119768

20. Hubaud, A., Regev, I., Mahadevan, L., & Pourquié, O. (2017). Excitable Dynamics and Yap-Dependent Mechanical Cues Drive the Segmentation Clock. Cell, 171(3), 668–682.e11. 10.1016/j.cell.2017.08.043

21. Isomura, A., Asanuma, D., & Kageyama, R. (2026). Synchronization of the segmentation clock using synthetic cell-cell signaling. Genes & development, 40(1-2), 124–141. 10.1101/gad.352538.124

22. Jiang, Y. J., Aerne, B. L., Smithers, L., Haddon, C., Ish-Horowicz, D., & Lewis, J. (2000). Notch signalling and the synchronization of the somite segmentation clock. Nature, 408(6811), 475–479. 10.1038/35044091

23. Kageyama, R., Masamizu, Y., & Niwa, Y. (2007). Oscillator mechanism of Notch pathway in the segmentation clock. Developmental dynamics : an official publication of the American Association of Anatomists, 236(6), 1403–1409. 10.1002/dvdy.21114

24. Koch, F., Scholze, M., Wittler, L., Schifferl, D., Sudheer, S., Grote, P., Timmermann, B., Macura, K., & Herrmann, B. G. (2017). Antagonistic Activities of Sox2 and Brachyury Control the Fate Choice of Neuro-Mesodermal Progenitors. Developmental cell, 42(5), 514–526.e7. 10.1016/j.devcel.2017.07.021

25. Laird, D. W. (2014). Connexin hemichannels, gap junctions and their unconventional roles in development, disease and regenerative medicine. Nature Reviews Molecular Cell Biology, 15(12), 789–803. 10.1038/nrm3876

26. Lauschke, V. M., Tsiairis, C. D., François, P., & Aulehla, A. (2013). Scaling of embryonic patterning based on phase-gradient encoding. Nature, 493(7430), 101–105. 10.1038/nature11804

27. Lewis J. (2003). Autoinhibition with transcriptional delay: a simple mechanism for the zebrafish somitogenesis oscillator. Current biology : CB, 13(16), 1398–1408. 10.1016/s0960-9822(03)00534-7

28. Lewis, J., Hanisch, A., & Holder, M. (2009). Notch signaling, the segmentation clock, and the patterning of vertebrate somites. Journal of biology, 8(4), 44. 10.1186/jbiol145

29. Meijer, W. H. M., & Sonnen, K. F. (2024). From signalling oscillations to somite formation. Current Opinion in Systems Biology, 39, 100520. 10.1016/j.coisb.2024.100520

30. Oates, A. C., Morelli, L. G., & Ares, S. (2012). Patterning embryos with oscillations: structure, function and dynamics of the vertebrate segmentation clock. *Development (Cambridge*, England), 139(4), 625– 639. 10.1242/dev.063735

31. Oginuma, M., Moncuquet, P., Xiong, F., Karoly, E., Chal, J., Guevorkian, K., & Pourquié, O. (2017). A Gradient of Glycolytic Activity Coordinates FGF and Wnt Signaling during Elongation of the Body Axis in Amniote Embryos. Developmental cell, 40(4), 342–353.e10. 10.1016/j.devcel.2017.02.001

32. Oginuma, M., Harima, Y., Tarazona, O. A., Diaz-Cuadros, M., Michaut, A., Ishitani, T., Xiong, F., & Pourquié, O. (2020). Intracellular pH controls WNT downstream of glycolysis in amniote embryos. Nature, 584(7819), 98–101. 10.1038/s41586-020-2428-0

33. Reaume, A. G., de Sousa, P. A., Kulkarni, S., Langille, B. L., Zhu, D., Davies, T. C., Juneja, S. C., Kidder, G. M., & Rossant, J. (1995). Cardiac malformation in neonatal mice lacking connexin43. Science, 267(5205), 1831–1834. 10.1126/science.7892609

34. Solan, J. L., & Lampe, P. D. (2009). Connexin43 phosphorylation: structural changes and biological effects. The Biochemical journal, 419(2), 261–272. 10.1042/BJ20082319

35. Sonnen, K. F., Lauschke, V. M., Uraji, J., Falk, H. J., Petersen, Y., Funk, M. C., Beaupeux, M., François, P., Merten, C. A., & Aulehla, A. (2018). Modulation of Phase Shift between Wnt and Notch Signaling Oscillations Controls Mesoderm Segmentation. Cell, 172(5), 1079–1090.e12. 10.1016/j.cell.2018.01.026

36. Steventon, B., & Martinez Arias, A. (2017). Evo-engineering and the cellular and molecular origins of the vertebrate spinal cord. Developmental biology, 432(1), 3–13. 10.1016/j.ydbio.2017.01.021

37. Ton, Q. V., & Iovine, M. K. (2013). Determining how defects in connexin43 cause skeletal disease. *Genesis (New York*, N.Y*. :* 2000), *51*(2), 75–82. 10.1002/dvg.22349

38. Tsakiridis, A., & Wilson, V. (2015). Assessing the bipotency of in vitro-derived neuromesodermal progenitors. F1000Research, 4, 100. 10.12688/f1000research.6345.2

39. Verrier, L., Davidson, L., Gierliński, M., Dady, A., & Storey, K. G. (2018). Neural differentiation, selection and transcriptomic profiling of human neuromesodermal progenitor-like cells *in vitro*. *Development (Cambridge*, England), 145(16), dev166215. 10.1242/dev.166215

40. Wang, N., De Vuyst, E., Ponsaerts, R., Boengler, K., Palacios-Prado, N., Wauman, J., Lai, C. P., De Bock, M., Decrock, E., Bol, M., Vinken, M., Rogiers, V., Tavernier, J., Evans, W. H., Naus, C. C., Bukauskas, F. F., Sipido, K. R., Heusch, G., Schulz, R., Bultynck, G., … Leybaert, L. (2013). Selective inhibition of Cx43 hemichannels by Gap19 and its impact on myocardial ischemia/reperfusion injury. Basic research in cardiology, 108(1), 309. 10.1007/s00395-012-0309-x

41. Wymeersch, F. J., Huang, Y., Blin, G., Cambray, N., Wilkie, R., Wong, F. C., & Wilson, V. (2016). Position-dependent plasticity of distinct progenitor types in the primitive streak. eLife, 5, e10042. 10.7554/eLife.10042

42. Yang, W., Lampe, P. D., Kensel-Hammes, P., Hesson, J., Ware, C. B., Crisa, L., & Cirulli, V. (2019). Connexin 43 Functions as a Positive Regulator of Stem Cell Differentiation into Definitive Endoderm and Pancreatic Progenitors. iScience, 19, 450–460. 10.1016/j.isci.2019.07.033

